# Dual and spatially resolved drought responses in the Arabidopsis leaf mesophyll revealed by single-cell transcriptomics

**DOI:** 10.1101/2024.08.30.610433

**Authors:** Rubén Tenorio Berrío, Eline Verhelst, Thomas Eekhout, Carolin Grones, Lieven De Veylder, Bert De Rybel, Marieke Dubois

**Affiliations:** Department of Plant Biotechnology and Bioinformatics, Ghent University, 9052 Ghent, Belgium; Center for Plant Systems Biology, VIB, 9052 Ghent, Belgium; VIB Single Cell Core, VIB, Ghent/Leuven, Belgium

**Keywords:** *Arabidopsis thaliana*, drought, leaf, transcriptome changes, single-cell RNA sequencing, mesophyll

## Abstract

Drought stress imposes severe challenges on agriculture by impacting crop performance. Understanding drought responses in plants at a cellular level is a crucial first step towards engineering improved drought resilience. However, the molecular responses to drought are complex as they depend on multiple factors including the severity of drought, the profiled organ, its developmental stage or even the cell types therein. Thus, deciphering the transcriptional responses to drought is specially challenging. Here, we investigated tissue-specific responses to mild drought in young *Arabidopsis thaliana* (Arabidopsis) leaves using single-cell RNA sequencing (scRNA-seq). To preserve transcriptional integrity during cell isolation, we inhibited RNA synthesis using the transcription inhibitor actinomycin D, demonstrating the benefits of transcriptome fixation for studying mild stress responses at single-cell level. We present a curated and validated single-cell atlas comprising 50,797 high-quality cells from almost all known cell types present in the leaf. We show that the mesophyll contains two spatially separated cell populations with distinct responses to drought: one enriched in canonical abscisic acid-related drought-responsive genes, and another one depicting iron starvation responses. Our study thus reveals a dual adaptive mechanism of the leaf mesophyll in response to mild drought and provides a valuable resource for future research on stress responses.

## INTRODUCTION

Drought is one of the most challenging stresses for agriculture, imposing significant limitations on crop production and yield (Boyer, 1982). Even when moderate, the decrease in water content initiates stress signaling cascades triggering adaptive mechanisms in root and shoot. The response and severity of the symptoms vary widely depending on the stress level and the developmental stage at which the stress occurs (Fuad-Hassan et al., 2008; Araus et al., 2012; Bledsoe et al., 2017; Verbraeken et al., 2021). Vegetative growth is a particularly vulnerable stage at which drought can severely affect plant performance and the timing of transition to the reproductive stage. To cope with drought, plants activate adaptive mechanisms, including changes in root architecture and an overall shoot biomass reduction. Limiting shoot growth reduces the evaporative surface of the plants and safeguards energy resources, which can be invested in drought resilience mechanism in case of life-threatening drought levels (Claeys and Inzé, 2013). As such, understanding plant responses to reduced water availability is essential to develop strategies that enhance plant performance and resilience under drought stress (Muller et al., 2011; Claeys and Inzé, 2013; Martignago et al., 2019; Dubois and Inzé, 2020; Simmons et al., 2021).

In the shoot, adaptive responses vary depending on the tissue, and even within the different cell types of a same tissue. In the leaf epidermis, stomatal closure is one of the first responses to maintain leaf water potential at the onset of drought (Bertolino et al., 2019; Laxa et al., 2019). In parallel, epidermal cell divisions and expansion of differentiating cells are limited to constrain leaf growth (Granier et al., 2006; Baerenfaller et al., 2012; Clauw et al., 2016; Dubois et al., 2017; Chen et al., 2021). Also in the inner leaf tissues, environmental stress responses are not homogenous: for example, vascular and mesophyll cells display different responses to abiotic stressors, such as UV stress (Berkowitz et al., 2021). In our previous study, we observed a trend for mesophyll cells to be strongly responsive to drought at the transcriptome level, but could not study specific responses in depth (Tenorio Berrio et al., 2022).

In the last decade, plant responses to moderate drought levels have been extensively studied using bulk transcriptome profiling (Harb et al., 2010; Wilkins et al., 2010; Baerenfaller et al., 2012; Ma et al., 2014; Clauw et al., 2016; Dubois et al., 2017), providing a broad overview of gene expression changes, yet, possibly overlooking the tissue-specific responses. Single-cell and single-nucleus RNA sequencing (sc/snRNA-seq) technologies provide powerful tools to dissect the unique transcriptomic signature of individual cell types or cell populations during plant development or during stress responses (recently reviewed in Nolan and Shahan, 2023; Zhu et al., 2023b; Grones et al., 2024; Tenorio Berrio and Dubois, 2024). Both scRNA-seq and snRNA-seq are routinely used methods, each with their own assets and drawbacks that are of importance in the study of stress responses. Generally, snRNA-seq is preferred because it allows to capture dynamic stress-responsive gene expression changes occurring in the nucleus and does not require enzymatic cell wall digestion, a process for which some organs or species are particularly recalcitrant (Grones et al., 2024). However, snRNA-seq also involves some drawbacks compared to scRNA-seq, being for instance a lower transcriptome coverage (Ding et al., 2020; Tian et al., 2020; Farmer et al., 2021; Kao et al., 2021; Conde et al., 2022; Neumann et al., 2022; Wang et al., 2023; Zheng et al., 2023). Additionally, snRNA-seq does not capture cytoplasmic and chloroplastic RNA, potential contributor to the leaf responses to stresses that impact chloroplastic processes. In contrast, scRNA-seq offers higher transcriptome coverage but requires enzymatic digestion of cell walls to isolate single cells in the form of protoplasts (Birnbaum et al., 2005; Wang et al., 2021; Liu et al., 2023b). Cell wall digestion affects gene expression, and can thereby introduce artifacts and biases in transcriptome analyses that might hinder the interpretation of mild responses upon treatments (Birnbaum et al., 2003; Wang et al., 2021). Fixing the transcriptional state of the samples prior to cell wall digestion has been proposed to overcome this issue (Procko et al., 2022; Grones et al., 2024; Wu et al., 2024). While transcriptome fixation is often used in scRNA-seq studies in the animal field (Attar et al., 2018; Wohnhaas et al., 2019; Sunkara et al., 2021; Pennitz et al., 2023), the possibility to apply it to plant samples to counteract the transcriptional response to cell wall digestion and thereby improve the understanding of plant single-cell responses to mild stress conditions, remains unexplored.

In this study, we dissected the tissue-specific responses to mild drought in young Arabidopsis leaves utilizing scRNA-seq. To preserve the drought-responsive transcriptional changes during cell wall digestion, we fixed the transcriptional state of the samples. For this, we applied actinomycin D (ActD), an anticancer drug blocking RNA synthesis by intercalating into DNA (Perry and Kelley, 1970; Sobell, 1985) and as such preventing additional transcriptional changes during cell isolation. This treatment improved the study of responses to mild drought, revealing responses that were largely masked or perturbed by cell isolation without transcriptome fixation. Our approach revealed two distinct drought-responsive mesophyll cell populations, one enriched in classical, abscisic acid-related drought-responsive genes, and another one associated with iron starvation. Using reporter lines and RNA fluorescence *in situ* hybridization (HCR-FISH), we located these two cell populations in two spatially separated regions of a drought-stressed young leaf, paving the way towards better understanding of the drought-adaptive mechanisms in Arabidopsis leaves.

## RESULTS

### Transcriptome fixation using actinomycin D preserves the transcriptional responses to mild drought

Because enzymatic digestion during protoplast generation from roots was shown to trigger strong transcriptional responses (Birnbaum et al., 2003; Wang et al., 2021), we speculated that the transcriptomic impact of cell wall digestion might mask the subtle responses to mild drought treatment, which were difficult to detect in our previous single-cell dataset (Tenorio Berrio et al., 2022). To circumvent this issue, we explored the use of transcriptome fixation during protoplast generation. Using bulk RNA-seq, we recorded the transcriptomic changes caused by the enzymatic cell wall digestion during protoplast generation to obtain a set of cell-wall digestion-responsive genes in the actively growing Arabidopsis leaves (third true leaf at 14 days after sowing, at this stage composed of dividing and differentiating cells, from well-watered (WW) conditions only) (**Figure 1A**). Enzymatic digestion during protoplast generation (for simplicity, the corresponding samples are named “Digested” samples) led to the differential expression of 11,041 genes (incl. 330 genes exclusively expressed in Digested samples) compared to the undigested full leaf (further named “Undigested” samples) used as a control (False Discovery Rate (FDR) < 0.05; or 6,707 genes with Log2 Fold Change (FC) >1, FDR < 0.01), representing around 55% of the 19,714 genes captured in the leaf transcriptome of the Undigested samples (**Figure 1B and Supplemental Table 1 and 2, Supplemental Figure 1A-B**). Subsequently, we explored the potential of ActD treatment to block the response to cell wall digestion by comparing the transcriptome of samples fixed during digestion (further named “Fixed-Digested” samples) with that of the Digested and Undigested samples. Four distinct gene expression profiles were identified when comparing the cell wall digestion-responsive genes in all three sample types (**Figure 1C, Supplemental Figure 1C, Supplemental Table 2**). Most of these genes displayed an attenuated induction (cluster 1; 6,101 genes) or repression (cluster 2; 4,965 genes) upon cell wall digestion when the samples were fixed (**Figure 1C**). Thus, transcriptome fixation during cell wall digestion can mitigate the transcriptomic changes due to protoplast generation in young Arabidopsis leaves.

**Figure 1.**
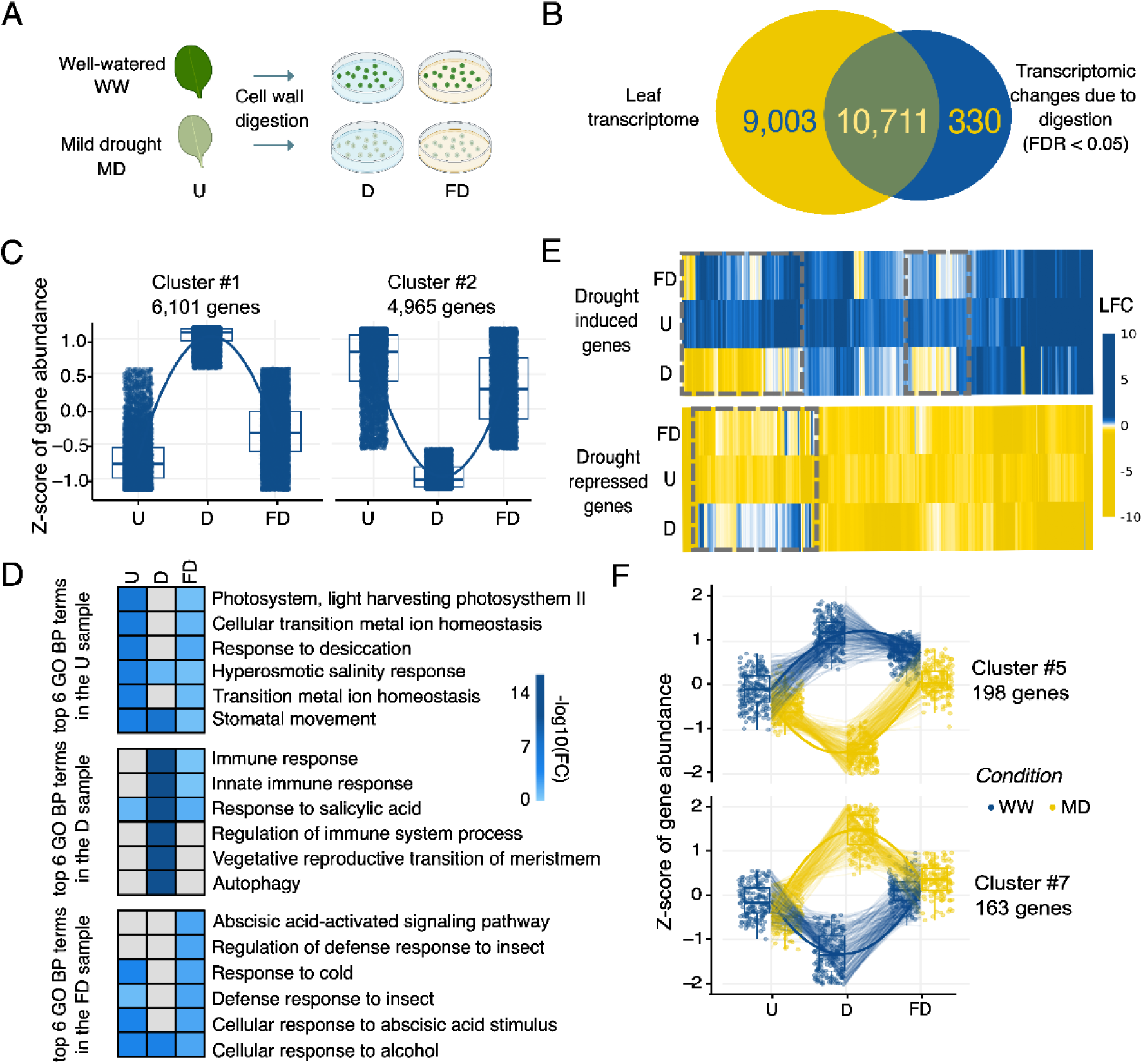
Cell-wall digestion responses in the leaf and interaction with the study of mild drought. (A) Schematic representation of the experimental workflow for bulk and scRNA-seq samples. Bulk RNA-seq was performed in the undigested leaf (U) sample (n=3) from plants grown under well-watered (WW) or mild drought (MD) conditions, and in samples that underwent cell wall digestion with or without fixation (digested leaf, D and fixed and digested leaf, FD) (n=2). scRNA-seq was performed with both D and FD samples (n=2). (B) Venn diagram depicting the transcriptomic impact of cell wall digestion (differential expression analysis WW, D vs. WW, U) compared to the leaf transcriptome (all transcripts present in WW, U). (C) k-means clustering performed on the differentially expressed genes (DEG) captured upon cell wall digestion. The two most abundant DEG clusters are shown (the rest of the clusters is shown in Supplemental Figure 1). For each sample, a boxplot represents the dispersion of the z-score of gene abundance. (D) Heatmap depicting the top-6 GO terms (biological processes) enriched in each sample and their level of enrichment in all samples. (E) Heatmap depicting, within the different samples, the behavior of the drought-responsive genes identified from the undigested leaf samples (701 genes). The top and bottom panel represent the log_2_ fold changes (LFC) of the genes up- and down-regulated in the U sample, respectively. (F) k-means clustering performed on the DEGs captured upon interaction analysis between the isolation method and growth condition. The two most abundant DEG clusters are shown (the rest of the clusters are shown in Supplemental Figure 3). For each sample, a boxplot represent the dispersion of the z-score of gene abundance in each plant growth condition, well-watered (WW) or mild drought (MD).

Next, to evaluate whether transcriptome fixation during the digestion process could preserve the transcriptional changes caused by drought stress, plants were grown on an automated watering platform (Skirycz et al., 2011) and were exposed to mild-drought (MD) or WW conditions (**Figure 1A**). A total of 701 genes (339 down- and 362 up-regulated; FDR < 0.05) were identified as being differentially expressed in the undigested leaves harvested during drought compared to the undigested leaves from WW plants (**Supplemental Figure 2A and Supplemental Table 3**). In samples undergoing enzymatic cell wall digestion, either without (Digested) or with fixation (Fixed-Digested), drought affected the expression of 2,990 and 286 genes, respectively (**Supplemental Figure 2B-C and Supplemental Table 3**). Gene Ontology (GO) analysis of the drought-responsive genes showed that biological processes typically associated with drought (e.g. ‘response to desiccation’) were enriched in the Undigested and Fixed-Digested samples, but not in the Digested samples (without fixation) (**Figure 1D**). These results thus suggest that transcriptome fixation allows to retain the drought response upon cell wall digestion of drought-treated plants.

Finally, we investigated the molecular interaction between drought and cell wall digestion in more detail. We studied the behavior of the 701 drought-responsive genes from Undigested leaves in the samples that underwent enzymatic cell wall digestion by comparing the fold changes upon drought. Surprisingly, Digested samples did not only show different degrees of mild drought responses, but even displayed completely opposite gene expression behaviors, compared to the Undigested samples (**Figure 1E**). The substantial disruption in the behavior of drought-responsive genes due to protoplast generation was again mitigated in the Fixed-Digested samples. Further investigation of the statistical interaction between cell wall digestion and drought responses identified up to 1,497 significantly interacting genes (P_Growth_Condition*Digestion_Treatment_ < 0.05) (**Figure 1F, Supplemental Figure 3 and Supplemental Table 4**). For most of these genes, fixing the transcriptome during cell wall digestion largely attenuated the interaction effect, without completely blocking it. Taken together, our bulk RNA-seq experiments suggest that ActD application during protoplast isolation reduces the post-harvest transcriptional changes and, thereby, allows to maintain the drought-triggered transcriptional responses in young leaves of drought-stressed plants.

### An optimized scRNA-seq atlas for mild drought responses in Arabidopsis leaves

To detect the tissue-specific transcriptional effects of mild drought in young Arabidopsis leaves, we sampled plants under WW and MD conditions for cell isolation either without (Digested) or with (Fixed-Digested) ActD treatment, in two independent experiments (**Figure 1A**). Protoplast suspensions were profiled by scRNA-seq using the BD Rhapsody microwell-based system. After quality and doublet rate control (**Supplemental Figure 4 and 5**), we combined data from all scRNA-seq samples, retaining a total of 152,793 high quality cells with a minimum Unique Molecular Identifier (UMI) count of 1,250 and a total of 26,150 expressed genes. We subsequently applied batch correction, reducing the batch effect between samples (see Methods) and performed unsupervised clustering (**Figure 2A**). From this combined dataset, 85,889 cells were isolated from Digested samples (48,072 WW and 37,817 MD), while 66,904 cells came from Fixed-Digested samples (38,641 WW and 28,263 MD). Although cells of WW and MD samples integrated well throughout the UMAP plot (**Figure 2B**), the main source of transcriptional variation was caused by the transcriptome fixation (**Figure 2C, Supplemental Figure 6**), supporting our conclusions drawn from the bulk RNA-sequencing (**Figure 1C**).

**Figure 2.**
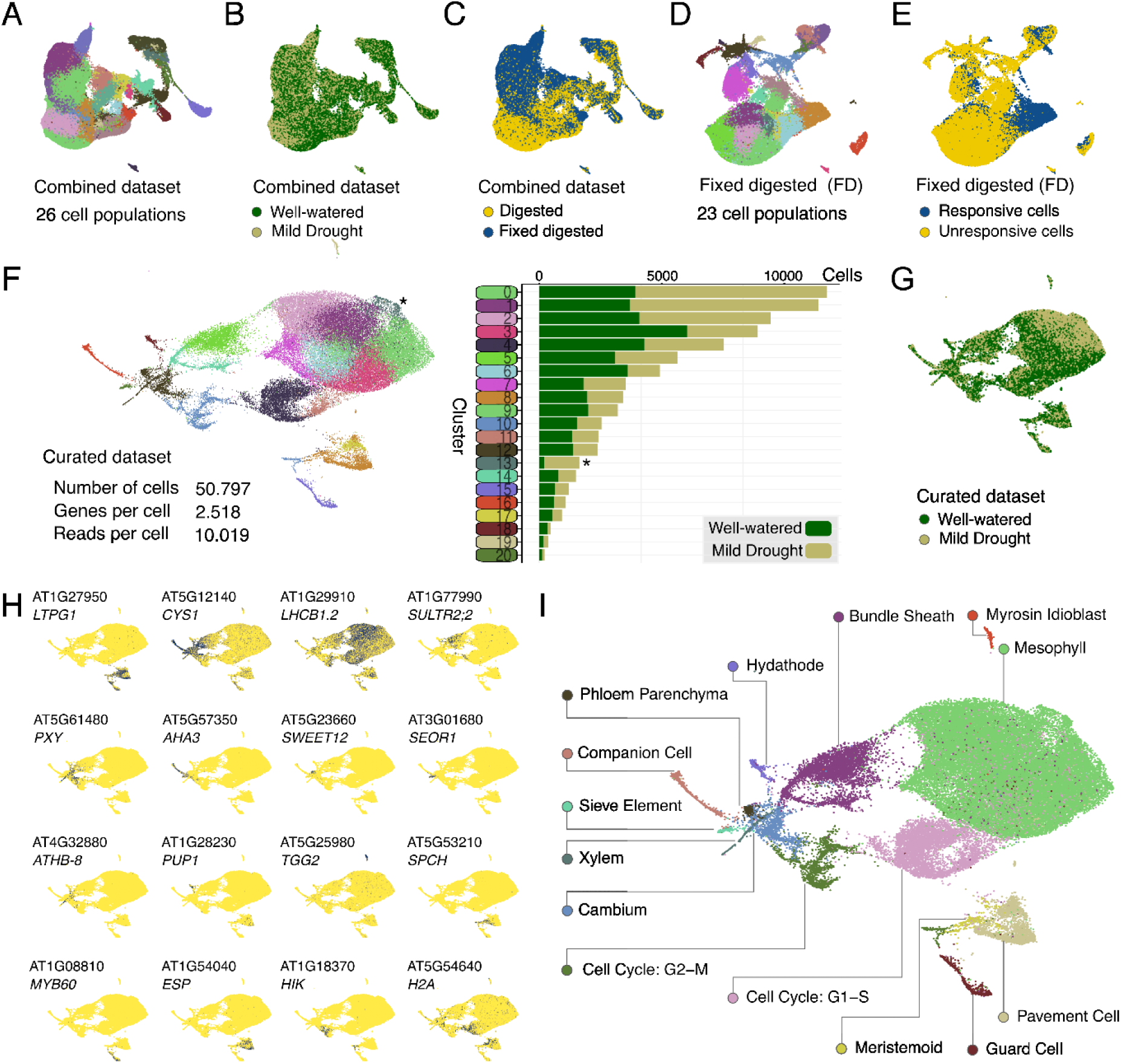
Optimizing and annotating a single-cell atlas for Arabidopsis leaves upon mild drought. (A-C) UMAP visualization of the combined dataset, containing both digested and fixed digested samples, following unsupervised clustering (A) or grouped by growth condition (B) or by fixation treatment (C). (D-E) UMAP visualization of the fixed digested samples, grouped by cluster (unsupervised clustering) (D) and by cell wall digestion response score, calculated based on the top-250 cell wall digestion-responsive genes (E). (F) UMAP visualization of the curated dataset containing cells of the Fixed-Digested samples with a low cell wall digestion response score. Cells are clustered by unsupervised clustering. The right panel represents the normalized number of cells per cluster grouped by growth condition. The asterisk highlights the drought-responsive “cluster 13”. (G) UMAP visualization of the curated dataset grouped by growth condition. (H) UMAP plots depicting the normalized expression of tissue-specific marker genes for the epidermis (*LPTG1*), vasculature (*CYS1*), mesophyll (*LHCB1.2*), bundle sheath (*SULTR2;2*), xylem (*PXY*), companion cells (*AHA3*), phloem parenchyma (*SWEET12*), sieve elements (*SEOR1*), cambium (*ATHB-8*), hydathode (*PUP1*), myrosin idioblast (*TGG2*), meristemoids (*SPCH*), guard cells (*MYB60*), pavement cells (*ESP*), and two populations depicting cell cycle states: G2M (*HIK*) or G1-S (*H2A*). (I) UMAP plot with the annotation of the curated dataset.

Having established the beneficial role of ActD treatment in preserving the drought responses in the leaf, we proceeded with the dataset of Fixed-Digested samples (**Figure 2D**). Based on the top-250 cell wall digestion-responsive genes (calculated comparing the bulk RNA-seq data from the Digested to the Undigested samples and excluding the genes interacting with the study of mild drought), we calculated a digestion-response score and visualized this score in the different datasets (**Supplemental Figure 7 and Supplemental Table 2**). We noticed that a fraction of cells in the Fixed-Digested dataset (24%; 16,107 out of 66,904), which were unevenly spread across the different cell populations, seemingly escaped the ActD treatment and still showed a strong cell wall digestion response (**Figure 2E and Supplemental Figure 7**). After removing those cells to curate the dataset, the optimized single-cell atlas of young leaves exposed to MD or WW conditions comprises 50,797 high quality cells, expressing a total of 23,717 genes (on average 2,518 genes), and grouped by unsupervised clustering into 20 distinct clusters (**Figure 2F and Supplemental Figure 8**). Nearly all clusters were composed of a similar proportion of cells from the WW and the MD samples, no stress-specific cell clusters were observed (**Figure 2 F,G**).

### Prediction and experimental validation of cell types

To predict the cluster annotations of the curated dataset obtained from fixed samples, we analyzed the expression of reported marker genes used for annotation in previous single-cell studies (**Figure 2H**) (Kim et al., 2021; Tenorio Berrio et al., 2022). This allowed us to classify the clusters into 14 distinct cell populations, including 12 cell types, and 2 cell state populations composed of cells of which the cell type signature is masked by a strong cell cycle state signature (**Figure 2I**). The cell type populations were further grouped by main tissue type (mesophyll, epidermis, and vasculature), and the cell cycle phase of all populations was predicted (**Supplemental Figure 9**). Following the same method, the datasets of Digested samples and the combined dataset (Digested + Fixed-Digested) were also annotated, allowing to compare the cell wall digestion response in the different leaf tissues (**Supplemental Figure 10 and Supplemental Table 5**). Although most of the cell-wall digestion response was shared across the three main tissues, tissue-specific responses were also present (**Supplemental Figure 11).**

To validate the predicted annotation of each population, we generated reporter lines for 25 genes with specific expression in most of the predicted cell types, by fusing the promoter region to the nuclear-localized green fluorescent protein (GFP) and β-glucuronidase (GUS) genes (**Table 1**, **Figure 3A and Supplemental Figure 12**). First, we confirmed via GUS staining that the expression pattern of these genes along the leaf corresponded to the expected main tissue (**Supplemental Figure 13**). Next, we selected one representative gene per tissue (**Figure 3A**) and analyzed their expression in more detail via confocal microscopy on cross-sections of leaves (**Figure 3B-C** and **Supplemental Figure 14**). In line with the predicted expression patterns (**Figure 3A** – top 3 rows), we found genes expressed in multiple cell types of one of the main tissues (vasculature, epidermis and mesophyll), although their expression levels could differ between the cell types of that tissue. For instance, the vascular marker gene *DJ1A* was expressed along most of the vasculature, with a higher expression in the phloem cells (**Figure 3C**). In addition, *EPIDERMAL PATTERNING FACTOR LIKE-9* (*EPFL9*) and *PROTODERMAL FACTOR1* (*PDF1*) marked the mesophyll and epidermal main tissues, respectively. Due to the photosynthetic nature of the bundle sheath cells, *EPFL9* was also found expressed along the bundle sheath (**Figure 3C**). Furthermore, we also generated and verified reporter lines for the bundle sheath (*CYTOCHROME P83A1* or *CYP83A1*), for most of the vascular cell types present in the leaf, including the cambium (*EXPANSIN 4* or *EXPA4*), the companion cells of the phloem (*PP2-A1*), the phloem parenchyma (*AT3G11930*), the xylem (*FASCICLIN-LIKE ARABINOGALACTAN-PROTEIN 12* or *FLA12*) and the myrosin idioblast (*TGG2*), and for the epidermal stomata (*AT1G04800*) and pavement cells (*AT5G63180*, here named *PAVEMENT CELL-RESTRICTED EXPRESSION*) (**Table 1**, **Figure 3C and Supplemental Figure 14**). Lastly, the reporter of *DIRIGENT PROTEIN 11* (*DIR11*) marked the hydathodes (**Figure 3A,C and Supplemental Figure 14**). Although no reporter line was generated for the sieve elements of the phloem nor for the meristemoids, the identity of the corresponding clusters could be validated based on the expression of the previously reported *SIEVE-ELEMENT-OCCLUSION-RELATED 1* (*SEOR1*) and *SPEECHLESS* (*SPCH*) marker genes, respectively (**Figure 2H**) (MacAlister et al., 2007; Pelissier et al., 2008). Taken together, all generated reporter lines confirmed the annotation of the cell clusters of the scRNA-seq atlas of the young Arabidopsis leaf. Based on the expression patterns of the markers validating each UMAP cluster, we translated the scRNA-seq data from our optimized dataset into a user-friendly tool for visualization of any gene expression in a browsable UMAP plot (http://www.single-cell.be/plants) and throughout a schematic Arabidopsis leaf section (**Supplemental Figure 15**, http://www.psb.ugent.be/shiny/trex.leaf/).

**Figure 3.**
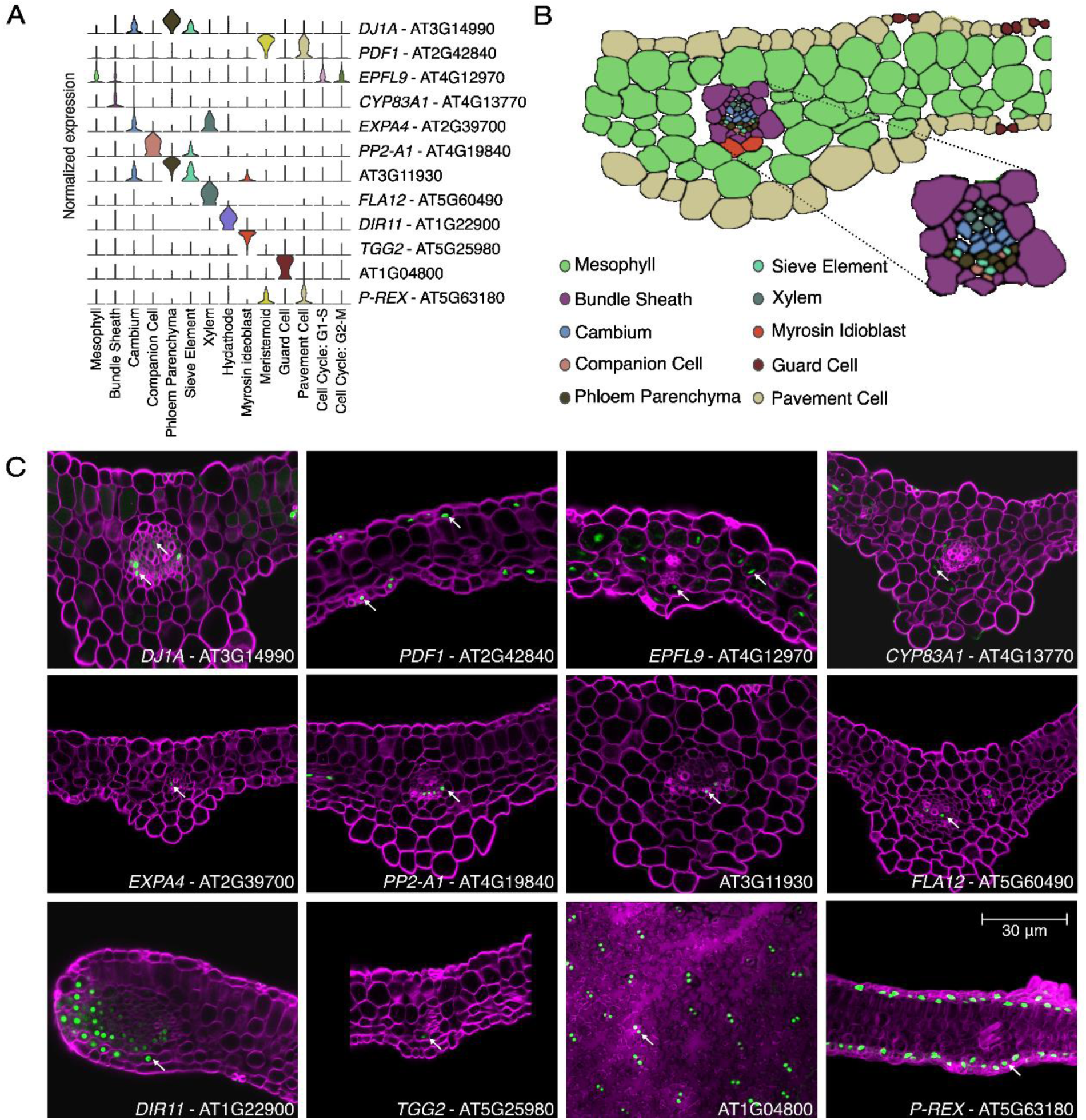
Biological validation of the annotation. (A) Violin plots depicting the tissue-specific expression of marker genes for the vasculature (*DJ1A*), epidermis (*PDF1*), mesophyll (*EPFL9*), bundle sheath (*CYP83A1*), cambium (*EXPA4*), companion cell (*PP2-A1*), phloem parenchyma (*AT3G11930*), xylem (*FLA12*), hydathode (*DIR11*), myrosin idioblast (*TGG2*), guard cell (*AT1G04800*) and pavement cell (*P-REX*), that were selected for the generation of reporter lines. (B) Schematic representation of the section of the leaf colored by cell type. (C) Confocal imaging of the selected reporter lines expressing the GFP-GUS reporter construct driven by the promoter of the indicated genes. For all imaging, except for the guard cell reporter line, cross-sections of cleared leaves were made to visualize the inner tissues. White arrows point to a GFP-positive nucleus. The magnification (scale bar = 30 μm) is the same in all panels.

**Table 1.** Library of tissue-specific reporter lines.

| Cell Type | Gene | Name |
| --- | --- | --- |
| Epidermis | AT2G42840 | *PDF1 – PROTODERMAL FACTOR1* |
|  | AT3G16370 | *APG* |
| Vasculature | AT1G64700 | Unknown gene |
|  | AT5G03610 | *GGL25* |
|  | AT1G80520 | Unknown gene |
|  | AT3G14990 | *DJ1A - DJ-1 HOMOLOG A* |
| Mesophyll | AT4G26530 | *FBA5 - FRUCTOSE-BISPHOSPHATE ALDOLASE 5* |
|  | AT2G34430 | *LHB1B1 - LIGHT-HARVESTING CHLOROPHYLL-PROTEIN COMPLEX II SUBUNIT B1* |
|  | AT4G12970 | *EPFL9 - EPIDERMAL PATTERNING FACTOR LIKE-9* |
| Bundle Sheath | AT1G25230 | Unknown gene |
|  | AT4G13770 | *CYP83A1 - CYTOCHROME P83A1* |
| Cambium | AT2G39700 | *EXPA4 - EXPANSIN 4* |
| Companion Cells | AT4G19840 | *PP2A1* |
|  | AT5G18600 | *GRXS2 - GLUTAREDOXIN 2* |
|  | AT1G64370 | Unknown gene |
| Phloem Parenchyma | AT3G11930 | Unknown gene |
|  | AT5G24800 | *BZIP9 - BASIC LEUCINE ZIPPER 9* |
| Xylem | AT3G10080 | Unknown gene |
|  | AT5G60490 | *FLA12 - FASCICLIN-LIKE ARABINOGALACTAN-PROTEIN 12* |
| Hydathode | AT1G22900 | *DIR11 - DIRIGENT PROTEIN 11* |
| Myrosin idioblast | AT5G26000 | *TGG1 - THIOGLUCOSIDE GLUCOHYDROLASE 1* |
|  | AT5G25980 | *TGG2 - THIOGLUCOSIDE GLUCOHYDROLASE 2* |
|  | AT3G16400 | *NSP1 - NITRILE SPECIFIER PROTEIN 1* |
| Guard Cells | AT1G04800 | Unknown gene |
| Pavement Cells | AT5G63180 | *PREX - PAVEMENT CELL-RESTRICTED EXPRESSION* |

### Mesophyll cells display two spatially distinct transcriptional responses to drought

Having validated the annotation of each cluster of the single-cell atlas allowed us to confidently profile the responses to mild drought per leaf tissue (**Supplemental Table 6**). For each cell type, we identified up- or downregulated drought-responsive genes and observed that a fraction of those genes was affected by drought in a tissue-specific manner (**Figure 4A**). Drought responses included GO biological terms associated with, for example, “Pectin biosynthesis” in the xylem, “Response to auxin” in the pavement cells, “Response to brassinosteroids” in the hydathodes, and “Response to salicylic acid” in bundle sheath and guard cells (**Supplemental Table 6**). Further analysis of the shared drought-induced genes highlighted similarities in the response between the mesophyll, the bundle sheath and the pavement cells (**Figure 4B**). On the contrary, the phloem populations, being the companion cells, sieve elements and phloem parenchyma, together with the guard cells, shared a large amount of genes downregulated under drought (**Figure 4B**).

**Figure 4.**
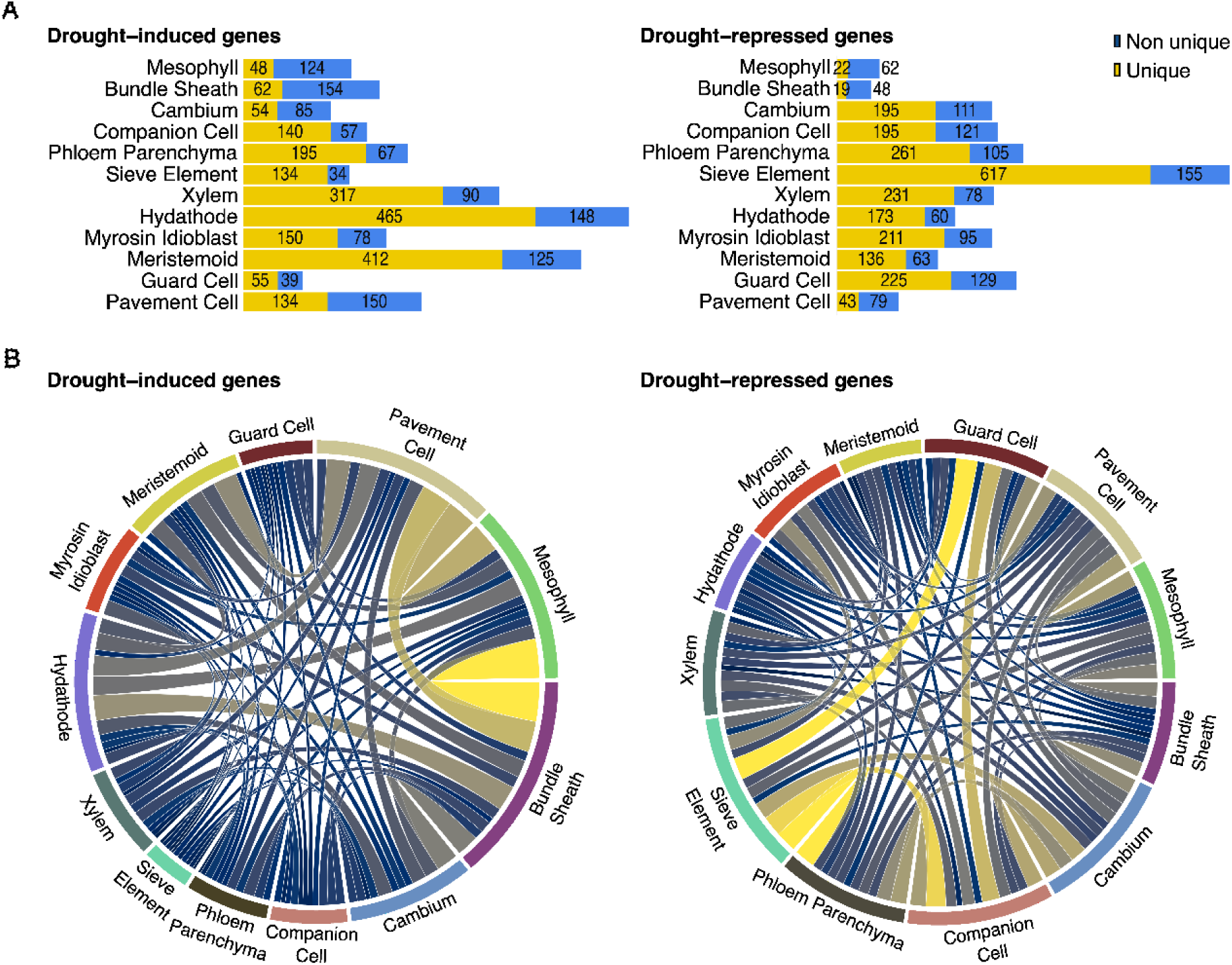
Shared and unique responses to mild drought. (A) Stacked bar plot displaying the number of uniquely and non-uniquely induced and -repressed genes for each cell type upon mild drought, in the left and right panel, respectively. (B) Chord diagram representing the relations between the shared drought-induced and -repressed responses between the different cell types, in the left and right panel, respectively. Connections are colored by the extent of the shared response, where dark blue and bright yellow connections represent few and large shared responses, respectively.

Although the mesophyll tissue did not appear to be the most responsive cell type based on the number of responsive genes, the mesophyll cluster “13” displayed a visibly higher percentage of drought-responsive cells after unsupervised clustering compared to any other cluster (**Figure 2F,I**). A closer inspection of the UMAP representation of mesophyll cells revealed a clear shift between the mesophyll cells from the WW samples vs. those from the MD samples, indicating outspoken transcriptome differences (**Figure 5A**). For example, the expression of the drought stress marker genes *AT14A-LIKE1* (*AFL1*), *RESPONSE TO DESICCATION 20* (*RD20*) and *FIBRILLIN1A* (*FIB*) followed this gradient of cells from WW and MD sample (**Figure 5B and Supplemental Figure 16A**) (Yang et al., 2006; Aubert et al., 2010; Kumar et al., 2015). Consistent with the bulk RNA-seq results and with our previous study (Tenorio Berrio et al., 2022), the drought-repressed gene *CUPPER SUPEROXIDASE 2* (*CSD2*) was found to be expressed across the opposite gradient (**Supplemental Figure 16B**). These expression patterns support the presence of a gradient in the mesophyll population as displayed in the UMAP plot, with an increasing drought response from the lower left corner (mainly WW cells) towards the upper right corner (mainly MD cells). Perpendicular to this gradient, a second gradient following the abaxial-adaxial leaf polarity axis was observed, reflected by the expression of the adaxial and abaxial mesophyll markers *PHENYLALANINE AMMONIA-LYASE 1* (*PAL1*) and *CORONATINE INDUCED 3 (CORI3*), respectively (**Figure 5C-D**) (Procko et al., 2022).

**Figure 5.**
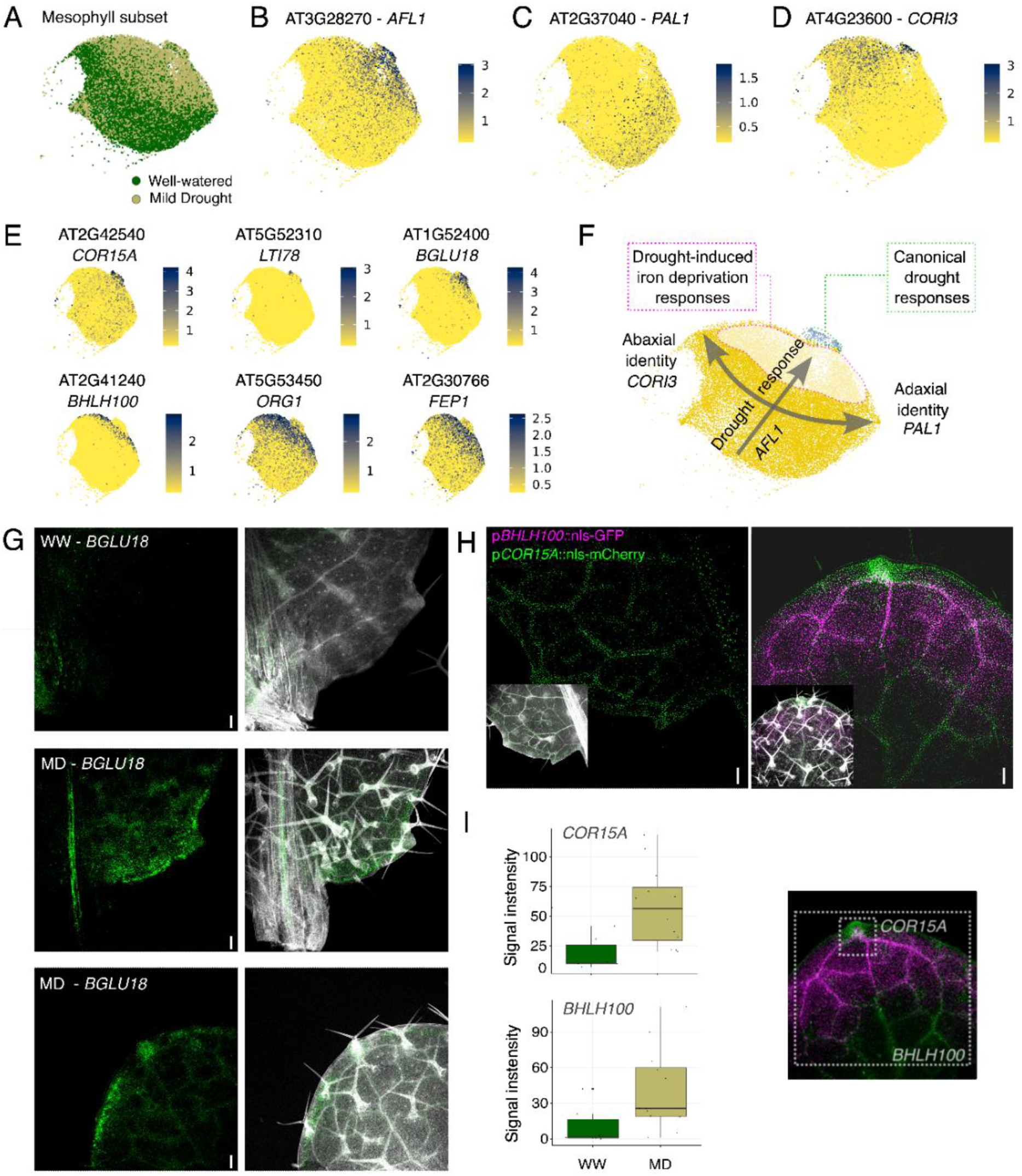
Mesophyll cells display two distinct drought responses. (A) UMAP visualization of the mesophyll subset of cells, grouped by growth condition. (B,C,D) UMAP plots depicting the normalized expression of markers of drought response (*AFL1*) (B), adaxial polarity (*PAL1*) (C) and abaxial polarity (*CORI3*) (D). (E) UMAP plots depicting the normalized expression of several drought-responsive genes, expressed either mainly in the tip zone of the mesophyll cluster (*COR15A*, *LTI78* and *BGLU18*), or in a large zone of the mesophyll cluster, but not in the very tip of the cluster (*BHLH100*, *ORG1* and *FEP1*). (F) Schematic representation of the two gradients present in the mesophyll populations: one gradient encompassing the drought response, starting in the zone with mainly well-watered sample cells, and ending in the upper right tip of the mesophyll cluster, and another gradient reflecting the leaf polarity. The highlighted parts represent the two drought-responsive cell populations. (G) Confocal microscopy of whole-mount fluorescence *in situ* hybridization (HCR-FISH) targeting the *BGLU18* transcripts, displaying the expression of *BGLU18* upon well-watered (WW) and mild drought (MD) conditions. Right images display white-stained cell walls of the leaf base (top and middle image) and tip (lower image). Scale bars = 100 μm. (H) Confocal imaging of GFP and mCherry expression in a dual *pbHLH100::nls-GFP* x *pCOR15A::nls-mCherry* reporter line exposed to mild drought conditions. Base (left) and tip (right) of the leaf, with the insets displaying white-stained cell walls of the corresponding zone. Scale bars = 100 μm. (I) Quantification of the fluorescent signal of the plant lines shown in (H), in the zones indicated in the picture, from leaves of plants grown under WW and MD conditions.

Despite some drought-responsive genes expressed commonly amongst the MD mesophyll cells (eg. *AFL1*), two distinct responses emerged within the group of drought-responsive cells (**Supplemental Table 7**). On the one hand, genes associated with canonical drought response processes like abscisic acid (ABA) metabolism, e.g. *BETA GLUCOSIDASE 18* (*BGLU18)*, the LEA protein *COLD-REGULATED 15A* (*COR15A*) or *LOW-TEMPERATURE-INDUCED 78* (*LTI78*), were expressed predominantly in the group of cells at the very tip of the drought gradient (**Figure 5E-F**, and other examples in **Supplemental Figure 16C)** (Hajela et al., 1990; Baker et al., 1994; Artus et al., 1996; Han et al., 2020). Notably, a population of cells undergoing such canonical drought response was not detected in the samples without fixation since, for example, the signature gene *BGLU18* was barely expressed in the Digested samples (**Supplemental Figure 17**). On the other hand, drought-responsive genes primarily associated with responses to iron deprivation or homeostasis, such as *BASIC HELIX-LOOP-HELIX PROTEIN 100* (*bHLH100*), *FE-UPTAKE-INDUCING PEPTIDE1* (*FEP1*) and *OBP3-RESPONSIVE GENE 1* (*ORG1*), were expressed in a large part of the MD mesophyll cells, but almost not in the cells at the very tip of the drought response gradient (**Figure 5E-F**, and other examples in **Supplemental Figure 16D)** (Wang et al., 2007; Sivitz et al., 2012; Hirayama et al., 2018; Kim et al., 2019). Consistently, this observation was supported by the enrichment of genes associated with responses to iron ion following an opposite gradient towards the WW-enriched mesophyll population, including FERRITINs (*FER1* and *FER4*) or *IRON SUPEROXIDASE DISMUTASE 1* (*FSD1*) (**Supplemental Figure 16B**) (Waters et al., 2012). Thus, we observed two gradients within the mesophyll cluster: a leaf polarity gradient and a drought-response gradient, with two distinct populations emerging under drought conditions (**Figure 5F**).

While the induction of genes like *BGLU18* and *COR15A* under drought was strong and detectable in the bulk RNA-seq profiling of the full leaf (**Supplemental Table 3**), the population of mesophyll cells displaying the canonical drought response was relatively small. To link the unique transcriptional profiles from the scRNA-seq analysis to a detailed spatiotemporal location within the leaf, we conducted whole-mount *in situ* hybridizations with probes targeting transcripts of *BGLU18* and *TONSOKU-ASSOCIATING PROTEIN 1* (*TSA1*), which was also specifically expressed in this population, in WW and drought-stressed leaves (**Figure 5E, Supplemental Figure 16**). Within the mesophyll, these genes were expressed in small patches of cells situated at the margin of leaves from drought-stressed plants, but not in well-watered samples (**Figure 5G and Supplemental Figure 18**). This was confirmed through cross-sections of the leaves, revealing additional expression in the phloem region of the midvein (**Supplemental Figure 18**).

Finally, we investigated whether the transcriptionally distinct large MD mesophyll cell population and the small cell population with the canonical drought response reflect two spatially separated mesophyll cell populations within a single leaf. We crossed two reporter lines to simultaneously visualize the expression of *bHLH100* and *COR15A*, two genes rarely co-expressed based on the scRNA-seq results (**Supplemental Figure 19**). *bHLH100* was highly expressed in the vasculature and mesophyll of the leaves, although only in a region towards the tip of the leaf (**Figure 5H**). Conversely, *COR15A* was expressed in mesophyll cells around the vasculature and at the margin of the leaf, aligning with the expression pattern of *BGLU18* (**Figure 5H**). Although the fluorescent reporters were also detectable in leaves of WW plants, the signal was increased in drought-stressed plants (**Figure 5I and Supplemental Figure 20**). In summary, the spatiotemporal resolution of the scRNA-seq dataset allowed us to identify two distinct drought-responsive mesophyll populations in the leaf: one at the margin and around the vasculature, and another encompassing the rest of the mesophyll cells. Although this observation mainly serves as a starting point for further functional studies on the role of these two cell populations during drought, this data reveals that the seemingly homogeneous mesophyll cells display a dual, early transcriptional response to cope with mild drought.

## DISCUSSION

In this study, we present a curated, comprehensive scRNA-seq atlas of young Arabidopsis leaves. Our atlas encompasses the transcriptome of leaves from well-watered and drought-stressed seedlings, capturing the vast majority of cell types present in the true leaf. However, some cell types could not be detected in our dataset and are also absent in previous leaf scRNA-seq atlases, probably due to technical constraints (Kim et al., 2021; Lopez-Anido et al., 2021; Procko et al., 2022; Tenorio Berrio et al., 2022; Zhu et al., 2023a). For example, trichomes could not be isolated, not through scRNA-seq, neither through snRNA-seq, possibly due to their low number or their large cell and nucleus size, resulting from high ploidy levels (Hulskamp et al., 1994; Churchman et al., 2006; Delannoy et al., 2023). Similarly, the very large glucosinolate-accumulating S-cells undergo programmed cell death at an early stage, and lack a known transcriptional signature, explaining why they are also unidentified (Koroleva et al., 2010; Kim et al., 2021; Procko et al., 2022; Tenorio Berrio et al., 2022; Delannoy et al., 2023; Maeda et al., 2023; Zhu et al., 2023a). In total, 12 distinct cell types and two populations of cells clustered by their cell cycle state were identified. Importantly, the annotation was further validated by reporter lines expressing *GFP-GUS* in each of these tissues of the young leaf. This atlas is now available as a browsable tool (http://www.single-cell.be/plants), along with a gene expression visualization tool (http://www.psb.ugent.be/shiny/trex.leaf/) in a schematic representation of the leaf.

Furthermore, our study explores the use of transcriptome fixation prior to scRNA-seq to preserve the transcriptomic impact of a subtle treatment, such as mild drought. Although widely applied in the animal field, tissue fixation in the plant scRNA-seq field is limited but holds great potential for limiting the impact of cell wall digestion on the plant’s transcriptome (Alles et al., 2017; Procko et al., 2022; Grones et al., 2024). Our bulk RNA-seq on digested samples (from well-watered conditions) revealed that more than half of the Arabidopsis leaf transcriptome is affected by cell wall digestion. Such massive transcriptomic changes are in line with a previous study that found ∼80% of the leaf transcriptome responding to overnight cell wall digestion (using the same statistical threshold as our study, FDR < 0.05) (Xu et al., 2021b). Thus, transcriptomic changes due to cell wall digestion in the leaf appear to be larger than in root tissues (Birnbaum et al., 2005; Chupeau et al., 2013). However, whether the leaf is more prone to digestion-triggered transcriptome reshuffling than roots, or whether other technical differences between these four studies could explain the differences between root and shoot, is not clear. In our study, although the cell wall digestion response was alleviated by transcriptome fixation, it was not completely inhibited. In fact, in the bulk RNA-seq analysis compared to the undigested leaf, most of the digestion-responsive genes were still de-regulated in the fixed samples. The scRNA-seq analysis of these samples suggested that there was no basal, general cell wall digestion response but that, instead, differences in the digestion response occurred between distinct cells. However, whether these differences are due to distinct effectiveness of fixation, or to differences in responsiveness to the cell extraction procedure, remains unknown. We speculate that this could be due to some cells being exposed longer to the digestion buffer before the fixative is added. Particularly relevant for our study, was the added value of transcriptome fixation to preserve the leaf response to mild drought, which appeared to be largely interacting with responses induced by the cell isolation procedure. This interaction could be caused, for example, by overlapping transcriptome changes or by drought-regulating proteins of which the synthesis is altered by ActD. Because of the extent of the drought and cell isolation interaction, it would be inappropriate to simply regress the cell wall digestion-responsive genes out of previous and future scRNA-seq stress studies; therefore, we recommend transcriptome fixation during protoplast isolation when aiming to compare subtle transcriptomic changes between treatments or genotypes. In summary, we proved the efficacy of transcriptome fixation to preserve stress response in single cells.

Our analysis also revealed two distinct mesophyll cell populations exhibiting contrasting responses to drought stress, a finding enabled by the transcriptome fixation prior to cell isolation. A first population, located at the margin of the leaf and around the vasculature, exhibits a typical drought response, consisting of genes induced by ABA, *COLD-REGULATED* (*COR*) genes and/or *EARLY-RESPONSIVE TO DEHYDRATION* (*ERD*) genes, such as *ERD12*, *KIN1*, *COR15A*, *COR47, BGLU18* and *LTI78* (Wang et al., 1995; Kim and Nam, 2010; Doner et al., 2021; Wu et al., 2023). We conducted whole-mount *in situ* hybridizations with probes targeting transcripts of *TSA1* and *BGLU18*, two of the most specific genes marking this population, to localize the cells with this typical drought response within the leaf. *BGLU18* is a β-glucosidase that hydrolyzes ABA-glucose ester (inactive form of ABA) to release free ABA in response to stress (Lee et al., 2006). *BGLU18* was previously shown to localize in leaf petioles, primarily in endoplasmic reticulum bodies, which are induced under stress, and to be involved in the early stages of ABA accumulation (Han et al., 2020). In fact, *TSA1*, which was reported to facilitate the formation of endoplasmic reticulum (ER) bodies in *Brassicales* (Geem et al., 2019), was also enriched in the leaf-margin mesophyll population. In addition, the observed expression pattern of *BGLU18* is consistent with the reported distribution of ER bodies in Arabidopsis rosette leaves (Nakazaki et al., 2019). Furthermore, under drought stress, BGLU18 was found to co-immunoprecipitate with the phloem-localized METACASPASE 3 (MC3), a positive regulator of drought response also transcriptionally inducing *BGLU18* (Pitsili et al., 2023). In addition, *LTI78* (also named *RESPONSE TO DESSICATION 29A, RD29A*), which is also enriched in this mesophyll population, is induced upon drought, high salinity and low temperature in Arabidopsis cotyledons (Nordin et al., 1993; Lee, 2010). *lti78* mutants were found to be more susceptible to drought, suggesting that LTI78 acts as a positive regulator of the drought stress response, avoiding plant death under severe drought (Liu et al., 2020; Liu et al., 2023a). Supporting this, *LTI78* expression is also upregulated by MC3, as well as by ABA, which was proposed to occur through the ERF transcription factor RAP2.6, a target of OPEN STOMATA1 (OST1) (Zhu et al., 2020; Ndathe et al., 2022; Pitsili et al., 2023). In this study, transcripts of the genes *TSA1* and *BGLU18*, which both co-expressed with *LTI78*, accumulated at the border of the leaf and in the phloem of the midvein upon drought. Similarly, the promoter of *COR15A*, which is also involved in ABA-dependent responses to environmental stresses and also targeted by RAP2.6 in the OST1 pathway, is induced upon drought at the border of the leaf and midvein (Gilmour et al., 1998; Gilmour et al., 2004; Zhu et al., 2020). In addition, we also observed *COR15A* expression in the mesophyll cells around the vasculature and at the tip of the leaf. Thus, *COR15A* is expressed in more widespread regions of the leaf than *BGLU18* and *TSA1*, but it remains unclear whether this broader expression is reflective of the wider expression partner displayed in our atlas or if it is influenced by the different techniques used for visualization (reporter line vs. *in situ* hybridization).

A second transcriptionally distinct mesophyll population responding to drought was localized in the more central, possibly distal, zone of the leaf. This population shared some drought responses with the population described above, but was characterized by an enrichment of genes associated to iron starvation. For example, multiple members of the basic helix-loop-helix (bHLH) transcription factor family, such as *bHLH38*, *bHLH39*, *bHLH100* and *bHLH101*, with roles in maintaining iron homeostasis, but also *FERRIC REDUCTION OXIDASE3* (*FRO3*) and *FEP1* genes were found enriched in this drought-induced population (Yuan et al., 2008; Sivitz et al., 2012; Kobayashi et al., 2014; Rasheed et al., 2016; Kurt and Filiz, 2018; Muhammad et al., 2022). Iron is an essential micronutrient for various cellular processes, including photosynthesis and respiration. Drought has previously been shown to impact iron homeostasis in sorghum roots, possibly to protect root cells against reactive oxygen species (Kim et al., 2019; Xu et al., 2021a). In addition, iron application has been shown to enhance drought tolerance in maize, canola and wheat (Tripathi et al., 2018; Rezayian et al., 2023; Abdullah et al., 2024; Mazhar et al., 2024). However, the precise molecular and functional connections between drought and iron homeostasis are still not well understood. It is possible that the reduction in transpiration in response to drought, resulting from decreased stomatal aperture and thickened leaf cuticula, limits the transport of nutrients, including iron, to the shoot, triggering an iron starvation response in the mesophyll cells.

In conclusion, we identified two mesophyll populations displaying different transcriptional responses to mild drought stress. We speculate that iron starvation responses first occur in the main photosynthetic cells that are more distant from the vasculature and, thus, from the micronutrient influx. Meanwhile, the mesophyll cells at the borders and midvein show early ABA responses, possibly serving as the first line of defense against drought or the initial recipients of water starvation signals from the root. The identification of two distinct mesophyll cell populations with different stress responses highlights the complexity of plant adaptation to drought, providing new insights into the tissue - or in this case within-tissue - specific responses underlying plant responses to mild drought.

## METHODS

### Plant material

Columbia-0 wild-type plants were used for all experiments. Reporter lines were generated by amplifying the 2kb genomic sequence upstream of the start codon (or up to the closest gene) for 25 selected marker genes (primers in **Supplemental Table 8**). The PCR fragment was cloned into pGGA000 via Gibson cloning, as described previously (Decaestecker et al., 2019). The following modules were assembled in pFASTRK-AG via a BsaI-mediated Golden Gate reaction: promoter in pGGA000, nls-GFP (without S) in pGGB000, GUS in pGGC000, 35S-terminator in pGGF000 with linkers where appropriate. After transformation, primary transformants were selected via the FAST-system and homozygous lines were obtained. A similar strategy was used for the generation of the COR15A reporter line. For the double reporter line (Figure 5), the pCOR15A::nls-mCherry line was crossed with pbHLH100::nls-GFP described in (Radoeva et al., 2019) and F1 progeny seeds were used.

### Growth conditions

Arabidopsis was grown on the weighing, imaging and watering automated machine (WIWAM) platform (http://www.wiwam.be) as described in our previous work (Tenorio Berrio et al., 2022). For transcriptomics, two experiments were performed fully independently from each other (∼6 months between).

### Sample preparation for bulk and scRNA-seq

For the bulk RNA-seq of undigested leaves, six third true leaves were pooled per condition, per biological replicate, immediately frozen in liquid nitrogen, and homogenized with the Retsch MM200. All other scRNA-seq samples were processed as follows. The cell wall digestion protocol was adapted from Ryu et al. (2019). Briefly, the third leaf of 80 plants per sample were harvested and chopped in 1 mL enzyme solution (0.4 M mannitol, 10 mM CaCl2, 20 mM MES, 20 mM KCl, 2% (wt/vol, 5.7 pH) cellulase, and 0.5% (wt/vol) macero enzyme). Chopped samples were transferred to a 70-μm cell strainer (model Corning, Merck) placed within a 6-well plate, in a total volume of 5 mL of the enzyme solution. In the fixed samples, 50 μM of actinomycin D (Sigma A9415) was added to the cell wall digestion solution. Samples were incubated, constantly shaking for 75 min at room temperature. After a 7 min centrifugation at 200 g, the pellet was resuspended in 255 uL washing solution (0.4 M mannitol, 10 mM CaCl2, 20 mM MES, and 20 mM KCl). Alive cells were enriched using magnetic levitation (LeviCell from Levitas Bio) by adding 45 μL of Levitation Agent to the cell suspension, followed by the separation of alive cells from dead cells and debris for 5 min. Subsequently, the purified cell suspension was retrieved from the LeviCell cartridge and counted using a Fast-Read® 102 counting slide. A total of 25,000 cells per sample were loaded in the Rhapsody (BD) or kept for bulk RNA-seq.

### RNA extraction, library preparation and RNA-seq analysis

Total RNA was isolated using the ReliaPrep RNA extraction kit (Promega) according to the manufacturer’s instructions. All samples were further processed by BGI TECH SOLUTIONS (Poland). The library was sequenced on the DNBseq platform, and around 30M 150 bp pair-end reads were aimed for. After initial filtering by BGI, the number of reads retrieved for each sample was around 20M. Subsequently, the data was processed using Galaxy. Reads were mapped on the Arabidopsis genome (TAIR 10) using the RNA-seq aligner tool STAR (Dobin et al., 2013). Statistical analysis was performed using the DESEQ package (Anders and Huber, 2010). DE analysis between two conditions/groups was performed using the Wald test. For more conditions/groups, the likelihood ratio test (LRT) was used. The false discovery rate (FDR) was controlled by adjusting P values using the Benjamini and Hochberg’s approach (Anders and Huber, 2010). DEGs were defined as genes with a FDR<0.05. The DEG distribution was mapped out using plots, which were generated in R using the EnhancedVolcano Package (Blighe K., 2019). The DEGs identified by each LRT were analyzed using the “DEGreport” package to identify patterns and generate clusters (Pantano, 2024). Z-score is the result of applying the scale() function, where values are centered to the mean and scaled to the standard deviation by each gene. A list of top cell wall digestion-responsive genes was obtained from the top 250 genes in cluster 1 of the cell wall digestion response, after removal of those cell wall digestion-responsive genes differentially expressed by drought (**Supplemental Table 2**).

### GO enrichment analysis

GO enrichment analysis was performed using the “clusterprofiler” package (Yu et al., 2012). Analysis was performed with the Benjamini & Hochberg’s adjustment method, with a q value cut-off of 0.4, and the results were simplified using the simplify() function. A less stringent threshold of 0.8 P-value was used for the tissue. The pheatmap package was used for visualization (Kolde, 2019).

### Single cell RNA-Seq sample preparation, library construction, and sequencing

For each sample, 25 000 cells were loaded on separate lanes of a BD HT Cartridge. Reverse transcription, cDNA amplification and library construction were performed following the manufacturer’s instructions (23-24117(02)). Libraries were sequenced on a NovaSeq 6000 flow cell (Illumina).

### Raw data processing and scRNA-seq analysis

The BD Rhapsody Sequence Analysis Pipeline (BD; version 2.0) was used to map the FASTQ files to the *Arabidopsis* reference genome (TAIR10). We utilized Seurat (V5.0.1) in R v4.16, to process and filter the expression matrix (Hao et al., 2024). Information and metadata parameters of the dataset are summarized in **Supplemental Table 9**. For each sample, low-abundance genes (all genes that were expressed in less than five cells) were removed. Cells expressing less than 1,000 genes or with less than 1,250 UMIs were filtered out. Normalization, detection of highly variable genes, scaling, clustering, and dimensionality reduction were performed using Seurat. The absence of a strong doublet-effect on the clustering was verified as the doublet score calculated with scds, with an average hybrid score lower than 0.25 for all samples (Bais and Kostka, 2020). Harmony (v0.1.0) (Korsunsky et al., 2019a) was used to integrate the data across repeats. The combined dataset is integrated both across repeat and fixation treatment. Uniform Manifold Approximation and Projection (UMAP) was performed using the top 32 harmony-adjusted principal components, followed by clustering using a resolution parameter of 0.8. A differential expression analysis was performed by the Wilcoxon rank sum test using the package Presto (Korsunsky et al., 2019b). A threshold of p-value < 0.05, log2 FC > 0.75, for genes expressed in at least a 5% of the cells in the population where it is induced, was set to calculate tissue-specific responses to mild drought, which were then visualized using the chordDiagram() function included in the “circlize” package. To calculate the tissue-specific responses to mild drought, a threshold was set at Upset plot was generated using the input from the pseudobulk analysis (pvalue < 0.05; log2FC >0 .5) using the ‘UpsetR” package. Scoring single cells was based on the approach described in (Tirosh et al., 2016). Pseudobulk expression profiles were generated by averaging the expression using the function AggregateExpression(). DESEQ2 was used to perform a pseudobulk differential expression analysis across different cell types or samples. Upset plot was generated using the input from the pseudobulk analysis (pvalue < 0.05; log2FC >0 .5) using the ‘UpsetR” package.

### GUS staining

Leaves were collected for β-glucuronidase (GUS) staining at 14 DAS. GUS staining was performed after fixation in 80% acetone. Leaves were incubated in GUS buffer (1 mM X-GlcA, 0.5% (v/v) Triton X-100, 1 mM EDTA, pH = 8, 0.5 mM K3Fe(CN)6, 0.5 mM K4Fe(CN)6, and 500 mM sodium phosphate buffer, pH = 7) at 37°C overnight and cleared for 1-2 days in 100% ethanol. After clearing, samples were embedded overnight in lactic acid and imaged with the Leica MZ16 binocular microscope equipped with a TOUPCAM camera.

### Leaf clearing, sectioning and confocal imaging

The leaves were fixed in fixative buffer (4% (w/v) paraformaldehyde in 1 x phosphate-buffered saline (PBS), supplemented with Triton 0.1% (v/v)), applying vacuum (4 x 30 min, mixed in between) to promote infiltration. After removing the fixative and performing three washing steps of 5 min in PBS, samples were incubated in the clearing solution Clearsee-alpha for 10 days (Kurihara et al., 2021). Clearsee-alpha solution was freshly made by adding sodium sulfite (50 mM) to the Clearsee solution and replaced every 2 or 3 days. For sectioning, samples were embedded in agar (3.5% m/v in PBS) and sectioned in 150 μ m sections using the Leica VT1200 S vibratome. Fixed samples were stained overnight with 0.1% (v/v) Renaissance SR2200 in ClearSee at room temperature in the dark for two days. Confocal imaging was performed using a Leica SP8X confocal microscope equipped with a Leica SuperK laser and a 40× (HC PL APO CS2, NA = 1.10) water immersion–corrected objective or a 10× (HC PL APO 10x/0.40 C92) dry objective.

### Fluorescence in situ hybridization

The method described by Huang et al. (2023) was used, with the following specifications or modifications. The hybridization probes targeting the *BGLU18* and *TSA1* transcripts were designed by and purchased from Molecular Instruments (lot numbers RTL889 and RTL890, respectively). HCR^TM^ amplification hairpins labeled with the 488 fluorophore, compatible with Renaissance SR2200, were purchased from the same provider. Samples were fixed in 5 mL FAA (7.44% formaldehyde, 5% glacial acetic acid, 50% ethanol in water) in 6-well plates by vacuum infiltration for 1h. Dehydration was performed by 10 min treatments with 70%, 90%, 100%, 100% ethanol followed by two 5 min washing steps with methanol. Rehydration was done by sequential washing with 75%, 50%, 25% methanol in DPBS-T water for 5 min per step, before being transferred to DPBS-T for 5 min. The leaves were subsequently incubated in a 10X dilution of the cell wall digestion stock (Huang et al., 2023) for 5 min, washed 5 min in DPSB-T, fixed again in FAA (formaldehyde, alcohol, acetic acid) for 15 min, and washed twice with DPBS-T for 5 min. A proteinase-K treatment was performed as described earlier (Huang et al., 2023). Samples were subsequently transferred to FAA for 30 min before being washed twice with DPBS-T. Next, samples were pre-hybridized by incubation in 30% probe hybridization buffer for 30 min at 37°C, and placed overnight at −20°C. Following this, the samples were transferred to 1.2 mL of 30% probe hybridization buffer containing 9.6 uL of each hybridization probe set (1 uM stock) and incubated overnight at 37°C. Subsequently, samples were washed twice for 30 min in 30% probe wash buffer at 37°C and twice for 10 min in 5x SSCT. Next, samples were pre-amplified for 30 min in amplification buffer. Hairpin solutions were prepared according to Huang et al. (2023), using 3 pmol of hairpin h1 and 3 pmol of hairpin h2. After overnight incubation with hairpins, excess hairpins were washed away by three washes (20 min each) in SSCT (saline sodium citrate buffer with 0.1% Tween-20) buffer. Finally, the samples were cleared in Clearsee-alpha as described above.

### Image analysis

Quantification of fluorescence intensity was performed using the FIJI-ImageJ software. In brief, we used the split colors tool on each confocal image. Next, the total intensity signal for each gene was calculated in a selected area (**Figure 5I**).

## Funding

Part of this work was funded by Ghent University (‘Bijzonder Onderzoeksfonds Methusalem Project’ no. BOF08/01M00408); the European Research Council (ERC CoG PIPELINES; 101043257 to B.D.R., E.V. and T.E.); VIB TechWatch Funding to C.G.; Marieke Dubois is a post-doctoral fellow of Flanders Research Foundation (FWO no. 12Q7923N).

## Author contributions

RTB and MD conceptualized the work. RTB, EV, TE, CG, LDV, BDR and MD conceived experiments. RTB, EV, TE and MD performed the experiments. RTB,TE and MD performed data analysis. TE, RTB and MD conceived the data visualization tool. BDR, LDV and MD supervised the project. RTB and MD drafted the manuscript which was further improved by all authors.

## Acknowledgements

The authors would like to thank the Cell cycle group for the stimulating working atmosphere, Dr. Dirk Inzé for supporting this project, Dr. Frank Van Breusegem, Dr. Patrick Willems and Robin Pottie for fruitful discussions, Dr. Dolf Weijers for kindly sharing seeds of the bHLH100 reporter line, and Dr. Annick Bleys for her help to improve the manuscript. We also would like to thank VIB Data Core for providing access and service to analyze our data on the Data Core servers. We are also thankful to the VIB Single Cell Core, VIB Flow Core Ghent and VIB Nucleomics for support and access to the instrument park (vib.be/technologies).

## Declaration of interests

The authors declare no competing interests.

## Data availability

Raw and processed data of the scRNA-seq experiments can be accessed at NCBI with respective GEO number GSE273033, together with the technical information following the guidelines of Grones et al., 2024 (**Supplemental Table 9**). The scRNA-seq data will be made accessible via an online browser tool upon acceptance of the manuscript (http://www.single-cell.be/plants). The raw bulk RNA-seq data can be accessed at NCBI with GEO number GSE273926.

## SUPPLEMENTAL INFORMATION

Supplemental Figure 1. Gene expression changes upon cell wall digestion.

Supplemental Figure 2. Gene expression changes upon mild drought.

Supplemental Figure 3. Expression profiles of genes with significant interaction between the growth condition and the cell isolation method.

Supplemental Figure 4. Quality control of the single-cell RNA sequencing samples.

Supplemental Figure 5. Droplet rate scores of the single-cell RNA sequencing samples.

Supplemental Figure 6. Transcriptional variation in the single-cell datasets.

Supplemental Figure 7. Cell wall digestion-response score.

Supplemental Figure 8. Quality control of the curated single-cell dataset.

Supplemental Figure 9. Main tissue populations and cell states in the curated dataset.

Supplemental Figure 10. Main tissue populations, cell states and annotation of remaining datasets.

Supplemental Figure 11. Tissue-specific and shared responses to cell wall digestion.

Supplemental Figure 12. Expression of additional tissue-specific marker genes.

Supplemental Figure 13. GUS staining images for the reporter lines of each tissue.

Supplemental Figure 14. Additional confocal microscopy images for the reporter lines of each tissue.

Supplemental Figure 15. Tool for single-cell data visualization in a leaf section.

Supplemental Figure 16. Expression of drought-related genes in the mesophyll tissue.

Supplemental Figure 17. Expression of *BGLU18* in the combined dataset.

Supplemental Figure 18. Transcript visualization of canonical drought responses.

Supplemental Figure 19. Co-expression of dual drought responses in the mesophyll.

Supplemental Figure 20. Microscopy images used for image analysis of the dual drought response.

Supplemental Table 1. Bulk RNA-seq: normalized data

Supplemental Table 2. Cell wall digestion response and gene clustering

Supplemental Table 3. Drought responses per sample type

Supplemental Table 4. Interactions between drought and cell wall digestion responses

Supplemental Table 5. Pseudobulk analysis of cell wall digestion responses

Supplemental Table 6. Drought responses per tissue and functional analysis

Supplemental Table 7. Drought-responsive genes in the leaf mesophyll

Supplemental Table 8. Primer list

Supplemental Table 9. Information and metadata parameters of the scRNA-seq dataset

## Notes

https://www.psb.ugent.be/shiny/trex.leaf/

http://www.single-cell.be/plants

### Competing Interest Statement

The authors have declared no competing interest.

