## Supplemental Figures for "Dual and spatially resolved drought responses in the Arabidopsis leaf mesophyll revealed by single-cell transcriptomics"

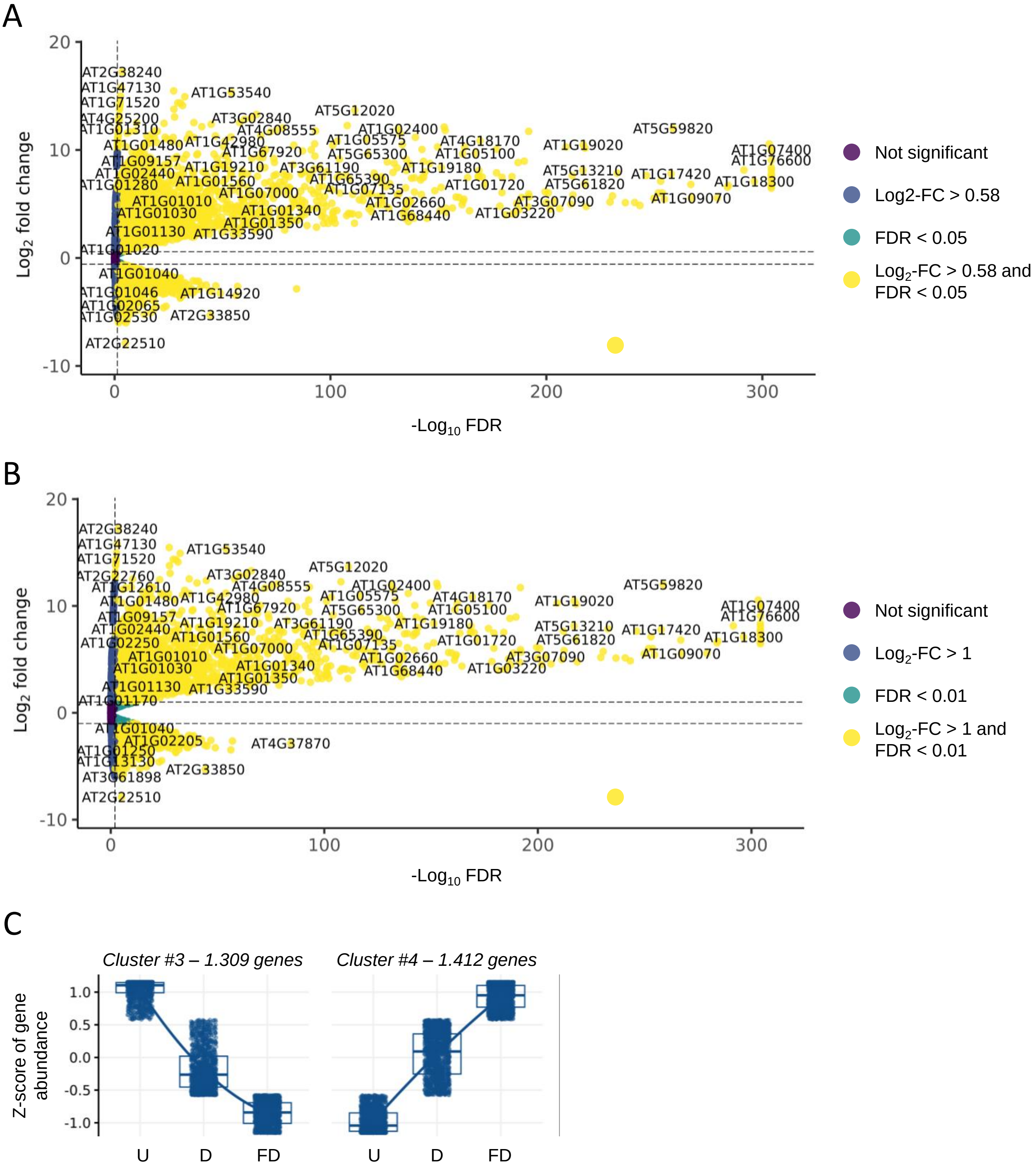


**Supp. Figure 1. Gene expression changes upon cell wall digestion. (A)** Volcano plot of the cell-wall digestion triggered response, using only False Discovery Rate (FDR) < 0.05 as a threshold for significance. **(B)** Volcano plot of the cell-wall digestion triggered response, using FDR < 0.01 and Log-2 Fold-Change (FC) > 1 as thresholds for significance. **(C)** k-means clustering performed on the differentially expressed genes captured upon cell-wall digestion without (D) and with (FD) fixation, compared to the undigested leaf tissue (U). The less abundant gene clusters are shown, while the most abundant clusters are shown in Figure 1C.

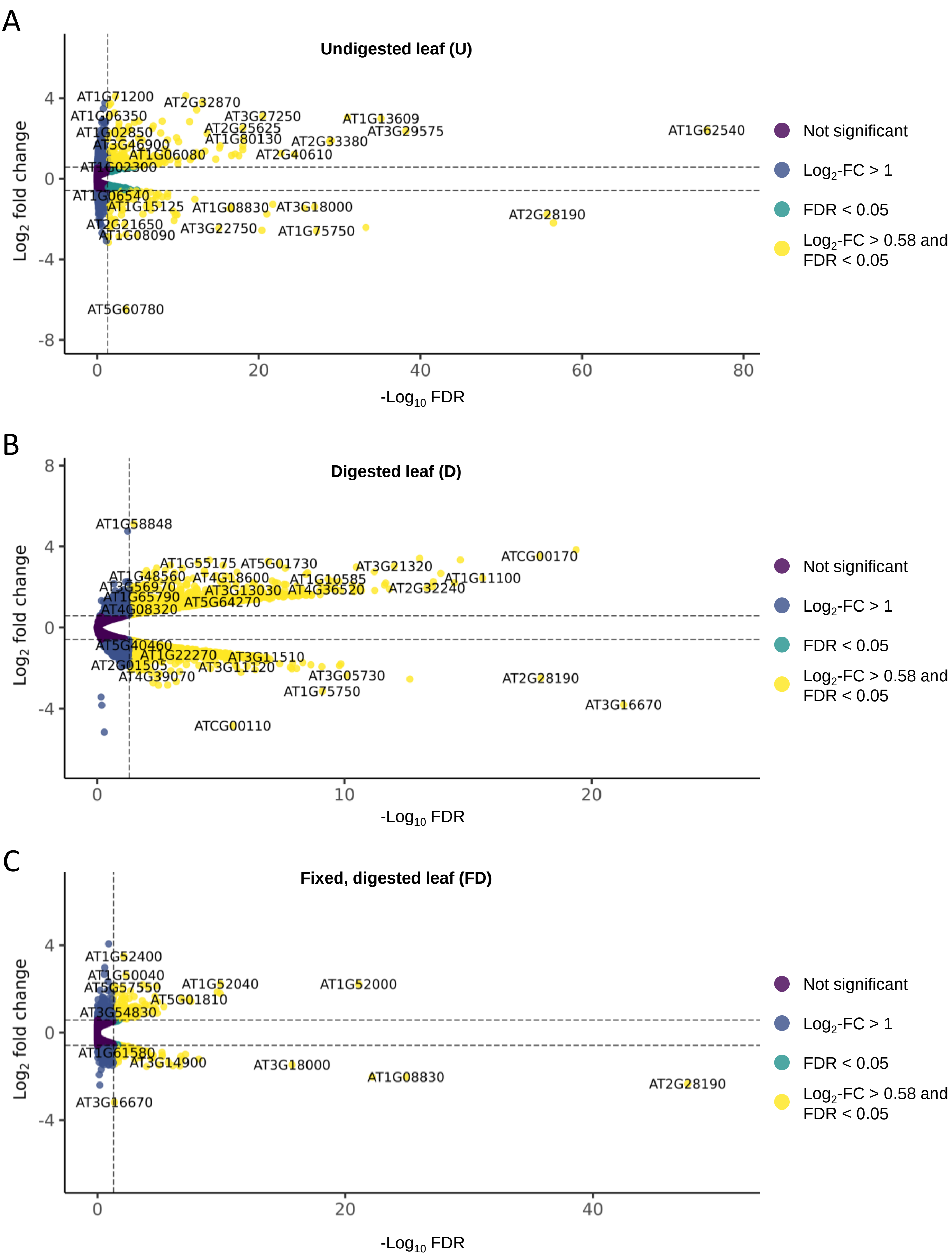


**Supp. Figure 2. Gene expression changes upon mild drought.** Volcano plots of drought-triggered response in undigested leaves **(A)**, digested leaves **(B)** and fixed an digested leaves **(C)**, using FDR < 0.05 and Log-2 FC > 1 as thresholds for significance.

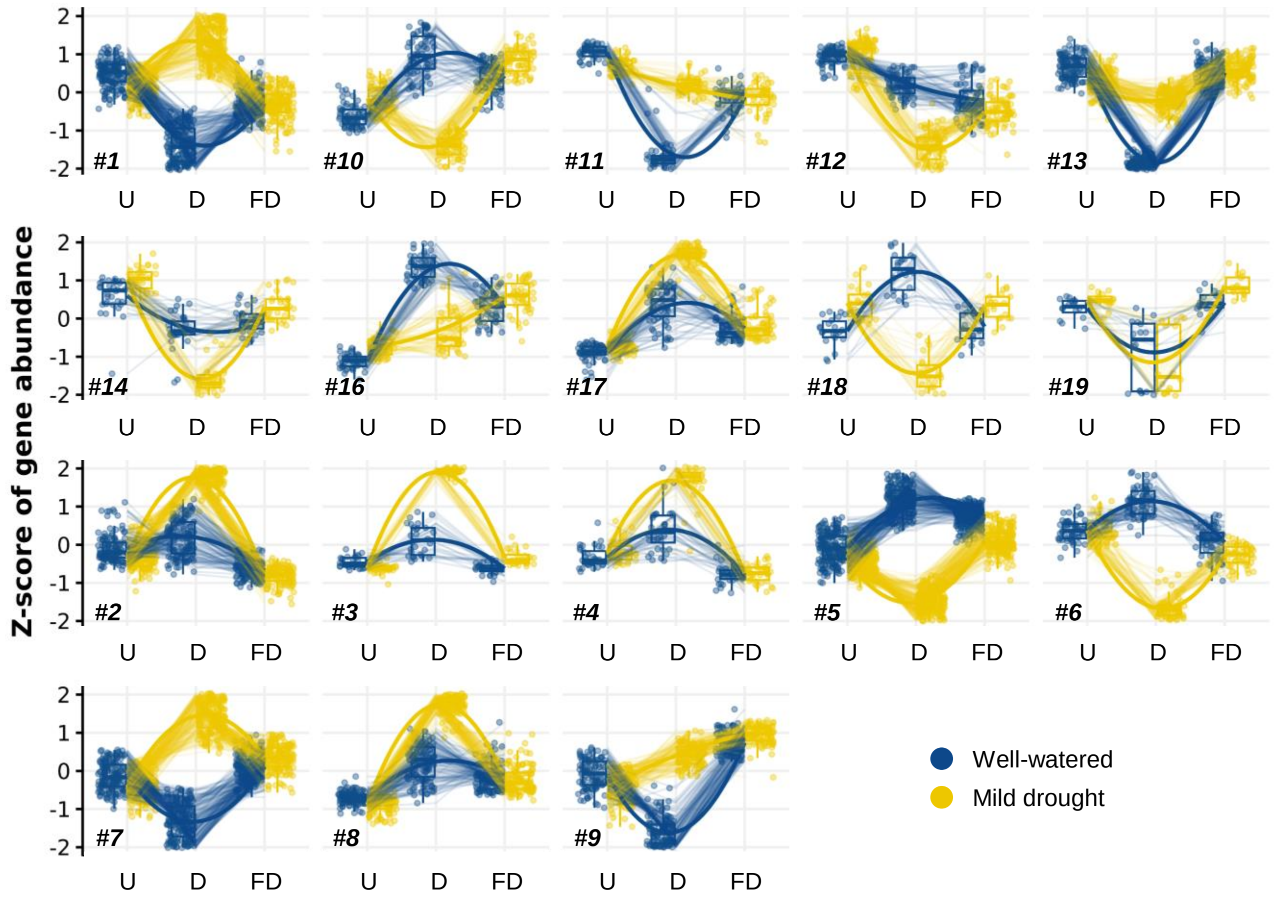


**Supp. Figure 3. Expression profiles of genes with significant interaction between the growth condition and the cell isolation method.** k-means clustering performed on the differentially expressed genes captured upon interaction analysis between the isolation method (digestion (D), fixation and digestion (FD), or undigested leaves (U)) and the growth condition. The number in the lower left corner indicates the cluster number. The list of genes within each cluster can be found in Supp. Table 4. For each sample, a boxplot represents the dispersion of the z-score of gene abundance in each plant growth condition, well-watered or mild drought.

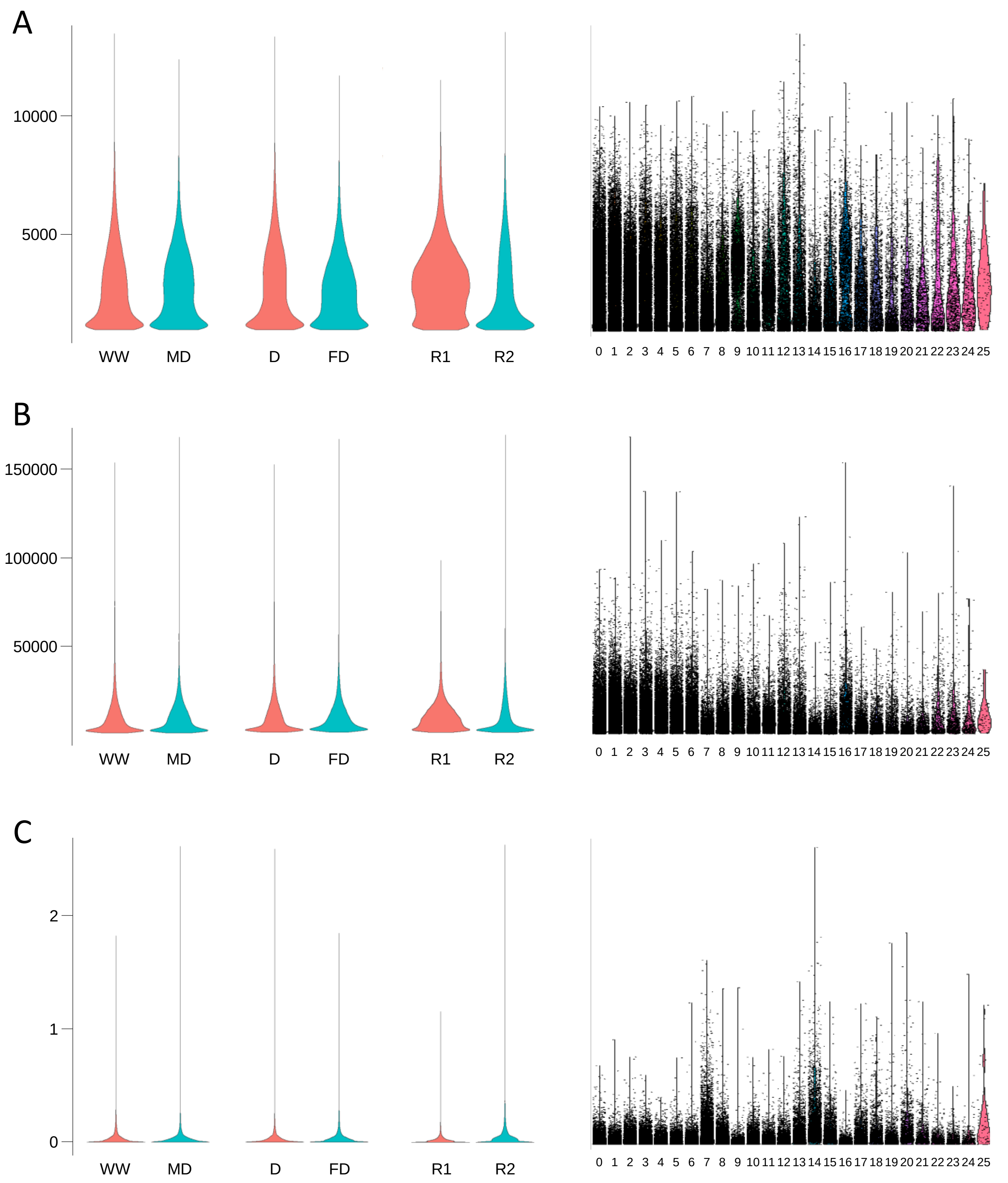


**Supp. Figure 4. Quality control of the scRNA-seq samples.** Violin plots representing the number of RNA features **(A)**, the number of counts **(B)** and the percentage of mitochondrial transcripts **(C)**. Data was grouped per condition (WW = well-watered, MD = mild drought), per cell isolation method (D = digested, FD = fixed and digested), per replicate (R = replicate) or per cluster (0-25).

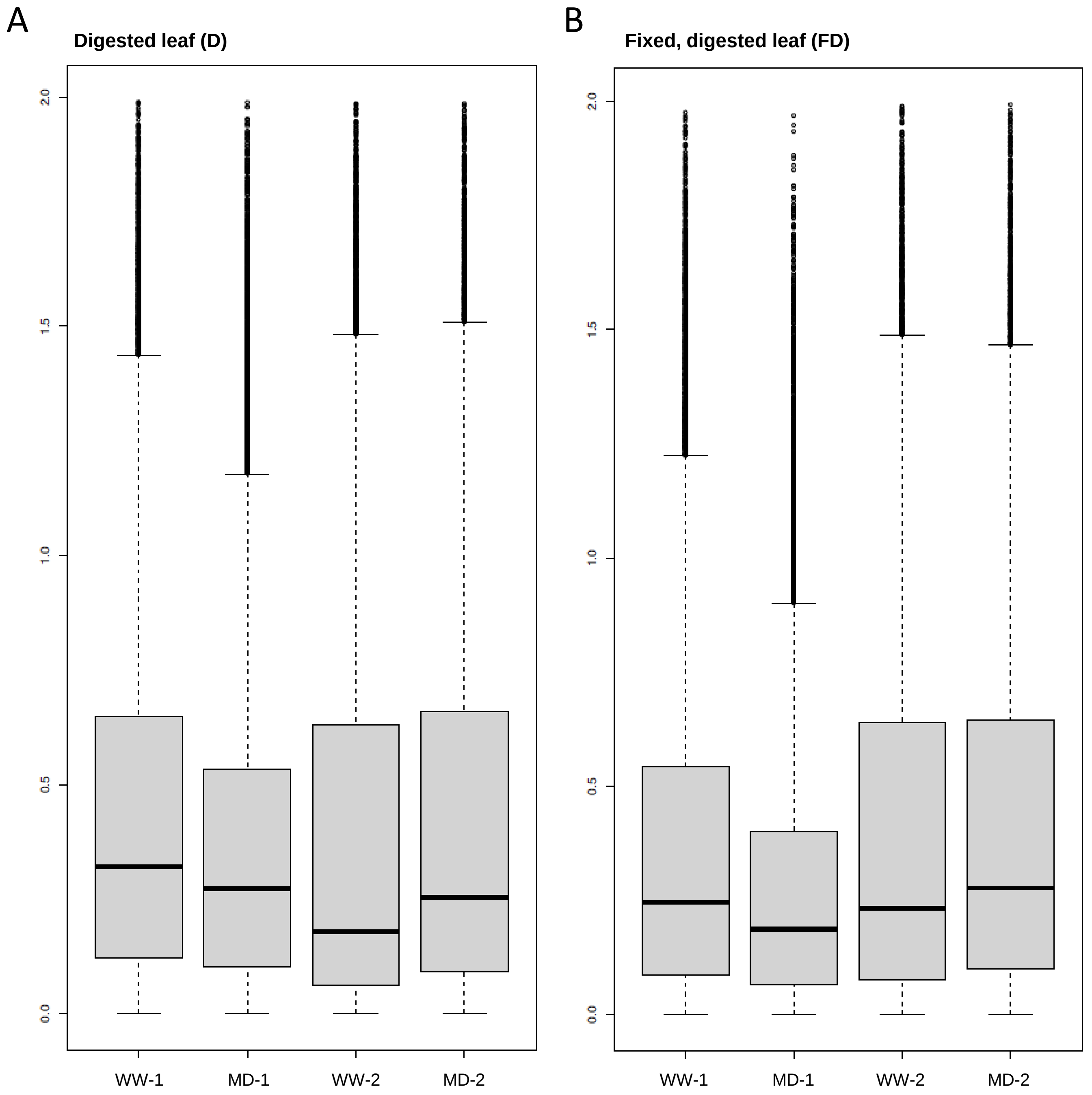


**Supp. Figure 5. Droplet rate scores of the scRNA-seq samples.** Boxplot depicting the hybrid droplet score, being a combination of the bcds and cxds methods, grouped by sample in the digested **(A)** and the fixed and digested datasets **(B)**.

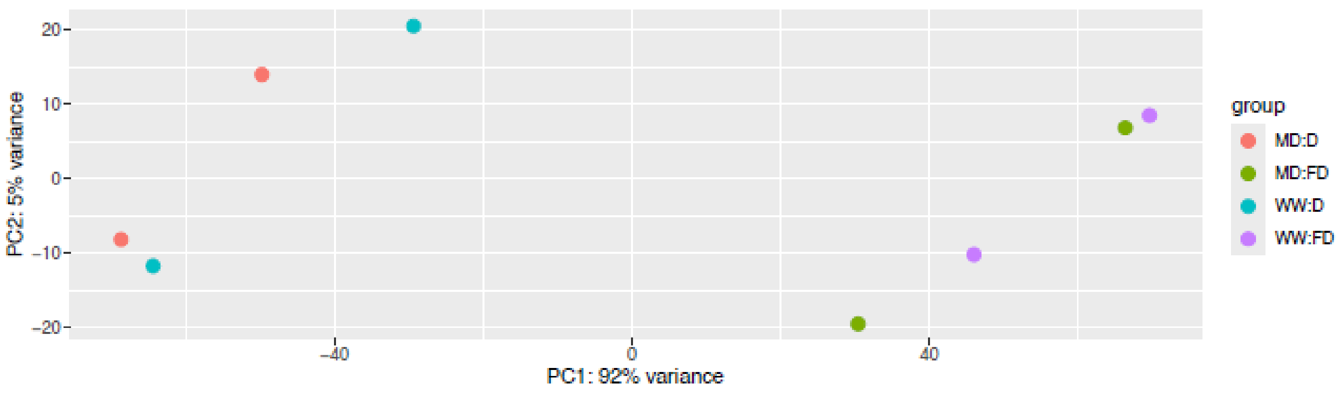


**Supp. Figure 6. Transcriptional variation in the single-cell datasets.** PCA plot depicting the two main principal components of the scRNA-seq samples.

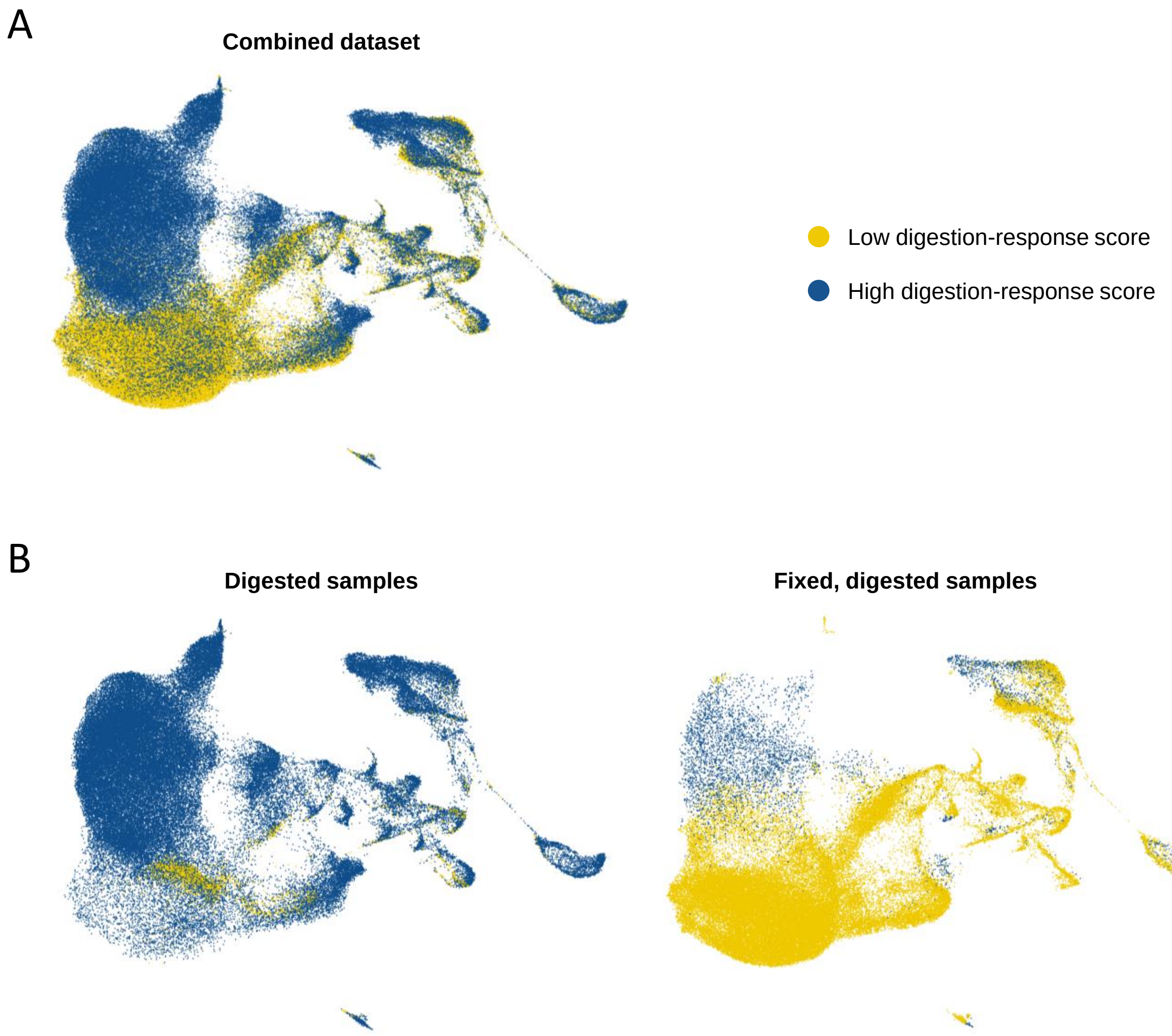


**Supp. Figure 7. Cell wall digestion-response score.** UMAP visualization of the combined dataset **(A)** and the dataset split by cell isolation method **(B)** with indication of cells showing a low or high cell wall digestion-response score. This score was calculated based on the top-250 cell wall digestion-responsive genes (calculated from the D samples, and excluding the genes interacting with the study of mild drought).

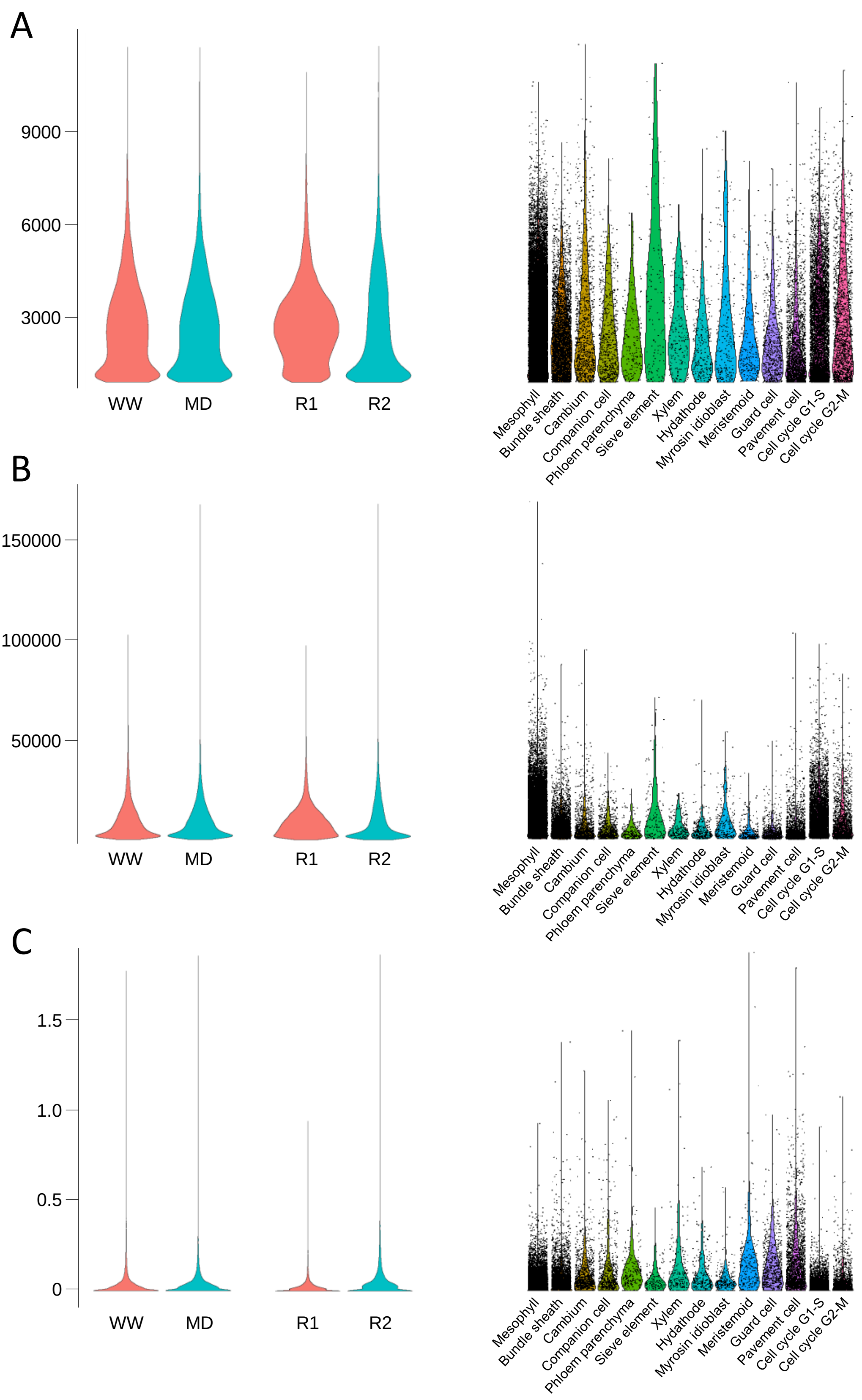


**Supp. Figure 8. Quality control of the curated single-cell dataset.** Violin plots representing the number of RNA features **(A)**, the number of counts **(B)** and the percentage of mitochondrial transcripts **(C)**. Data was grouped per condition (WW = well-watered, MD = mild drought), per replicate (R = replicate) or per cell type.

A

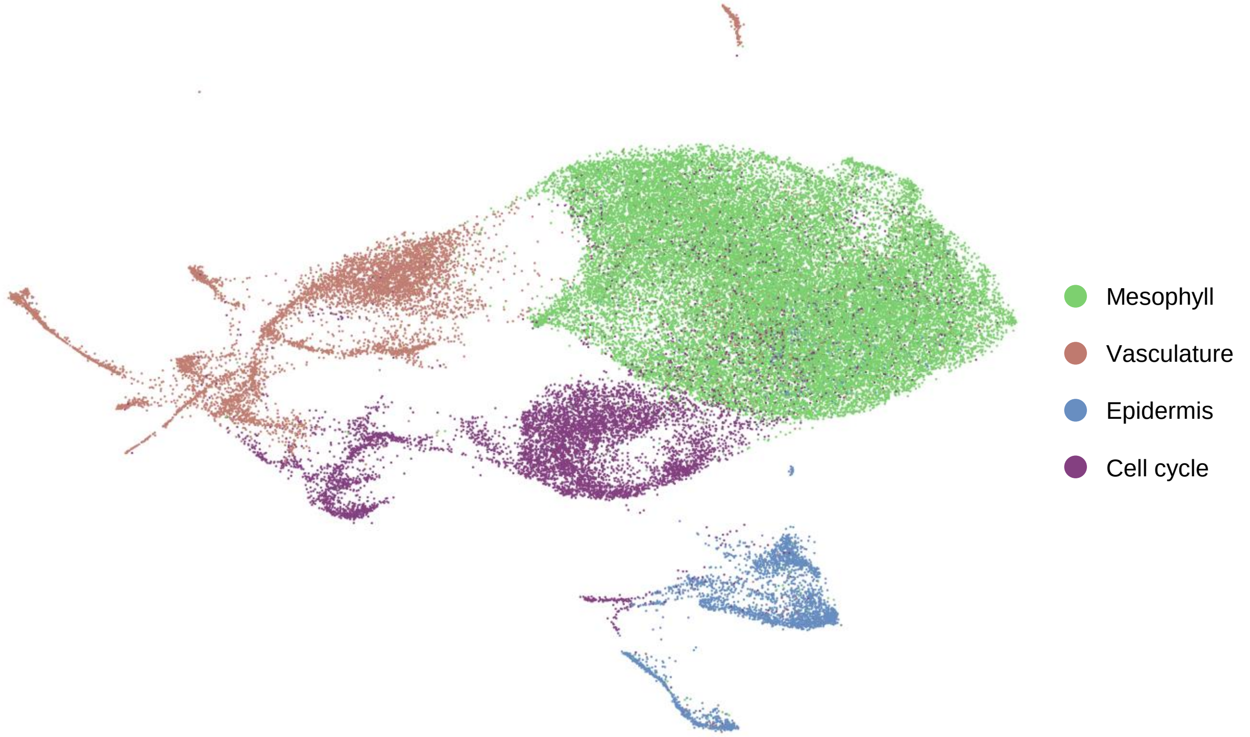


B

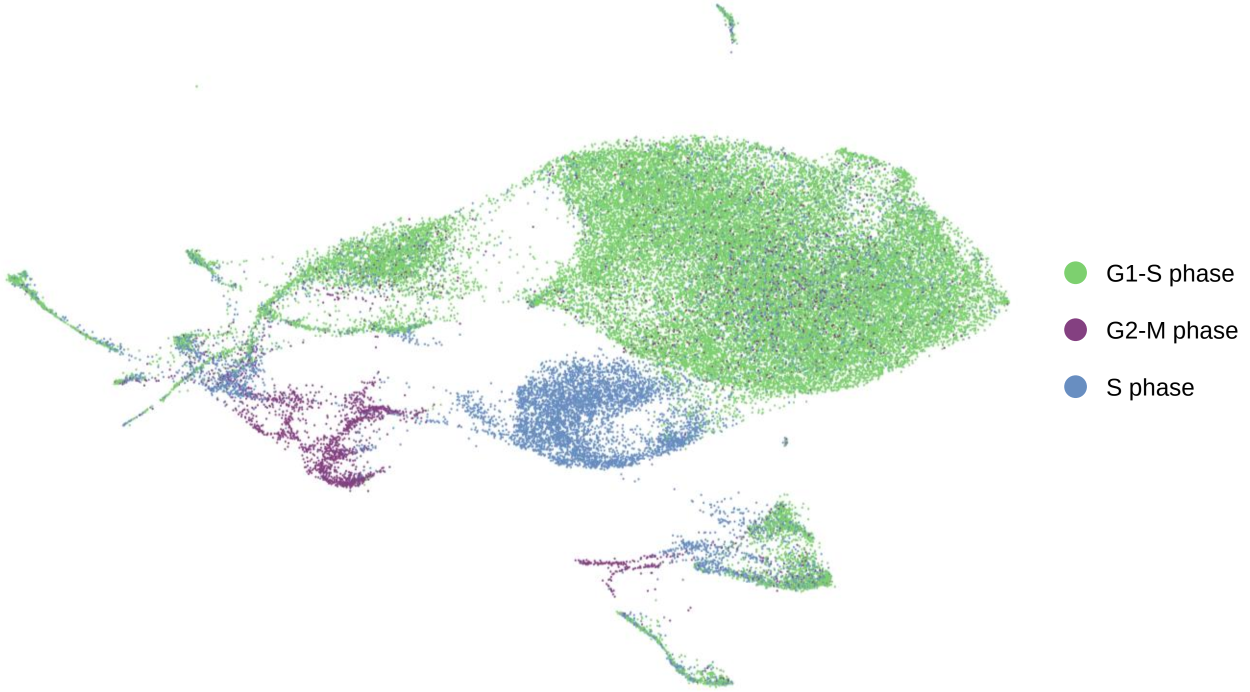


**Supp. Figure 9. Main tissue populations and cell states in the curated dataset.** UMAP plots depicting the curated dataset of fixed and digested samples, colored according to main tissue **(A)** or to cell cycle phase **(B)**.

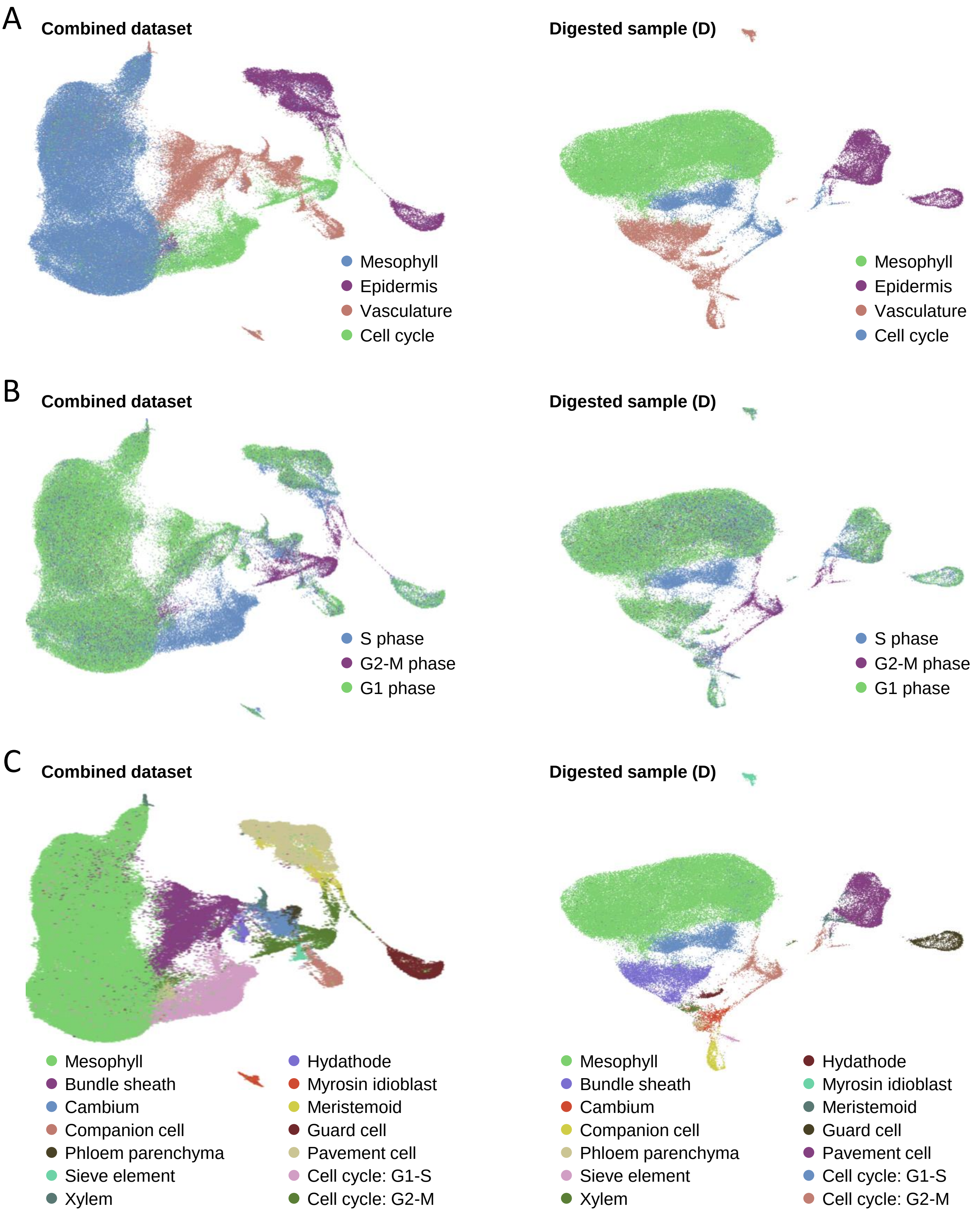


**Supp. Figure 10. Main tissue populations, cell states and annotation of remaining datasets.** UMAP plots depicting the combined dataset (left panels) and the one of digested samples without fixation (right panels), colored according to main tissue **(A)**, to cell cycle phase **(B)** or to cell type/state **(C)**.

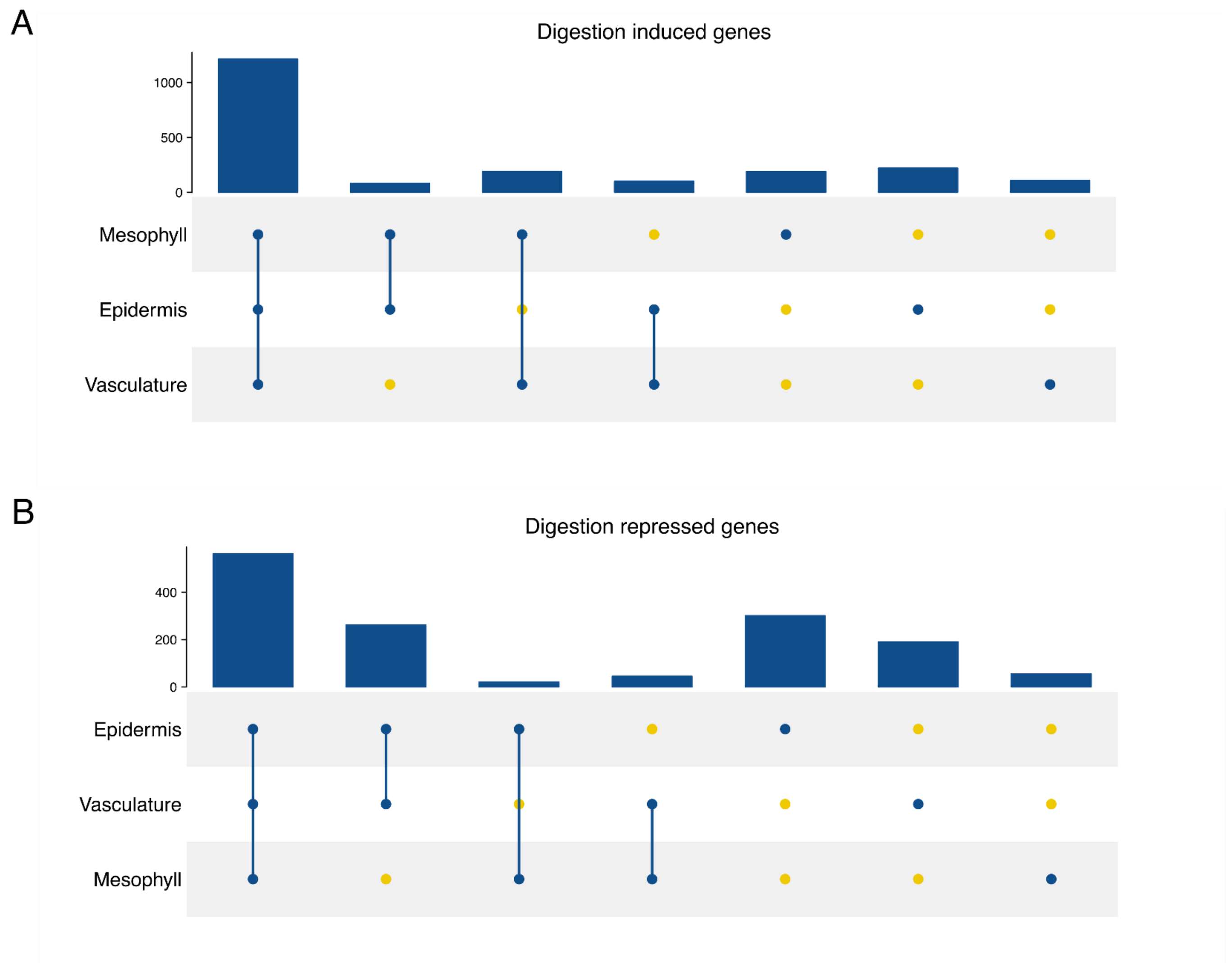


**Supp. Figure 11. Tissue-specific and shared responses to cell wall digestion.** Upset plot visualization of the total number of common genes among the selected main-tissues, depicted in the lower panel, between the genes induced **(A)** or repressed **(B)** upon cell wall digestion. Bar plots in the upper panel depicts the sizes of the intersections indicated in the lower panel.

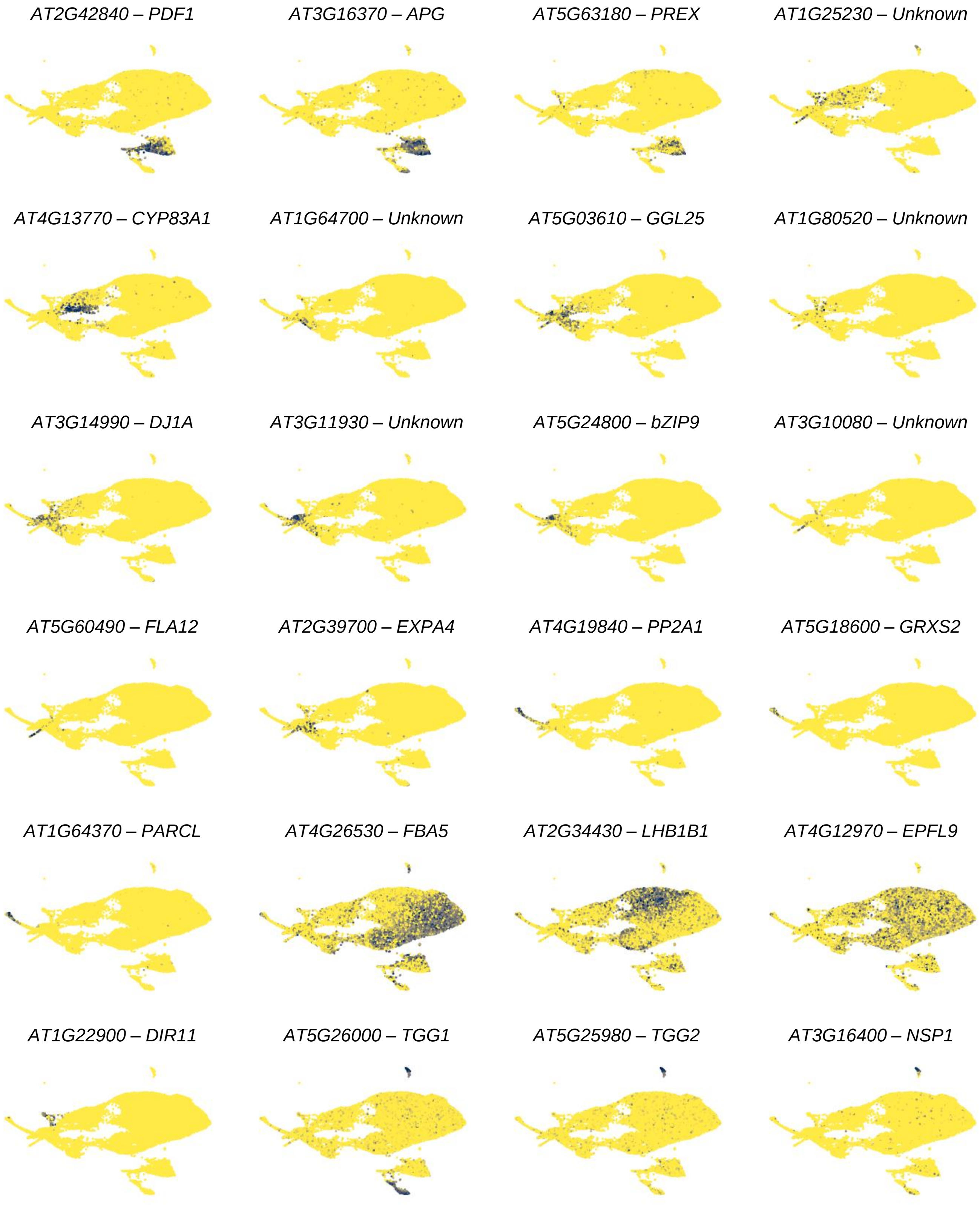


**Supp. Figure 12. Expression of additional tissue-specific marker genes.** UMAP plots depicting the normalized expression of additional marker genes for each tissue: epidermis (*PDF1*, *APG* ), pavement cell (PREX), bundle sheath (AT1G25230, *CYP83A1*), vasculature (AT1G46700, *GGL25*, AT1G80520 and *DJ1A*), phloem parenchyma (AT3G11930, *bZIP9*), xylem (AT3G10080, FLA12), cambium (EXPA4), phloem companion cell (*PP2A1*, *GRXS2*, *PARCL*), mesophyll (*FBA5*, *LHB1B1* and *EPFL9*), hydathode (*DIR11*), myrosin idioblast (*TGG1*, *TGG2* and *NSP1*).

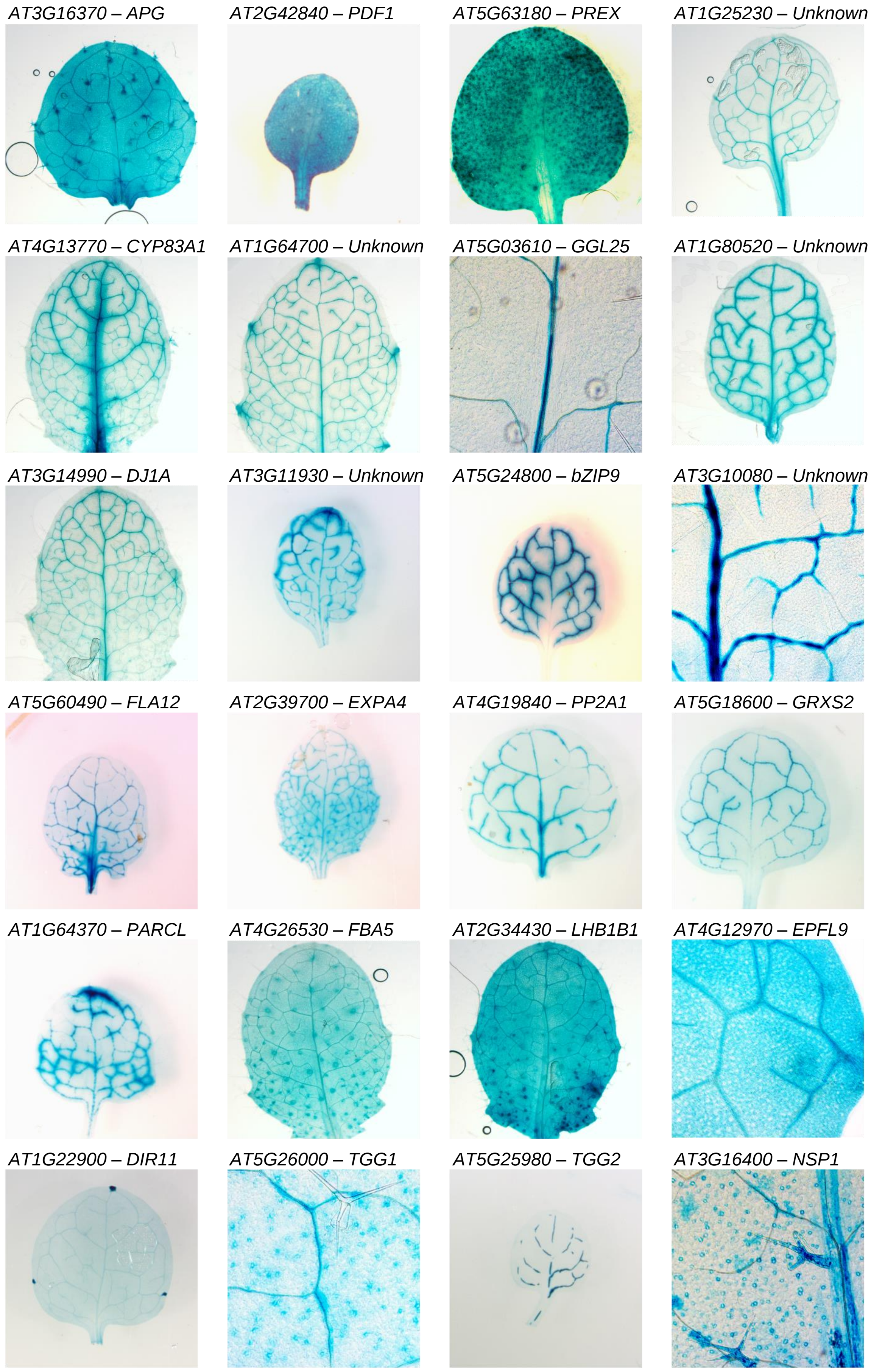


**Supp. Figure 13. GUS staining images for the reporter lines of each tissue.** Top-view images of true leaves are shown.

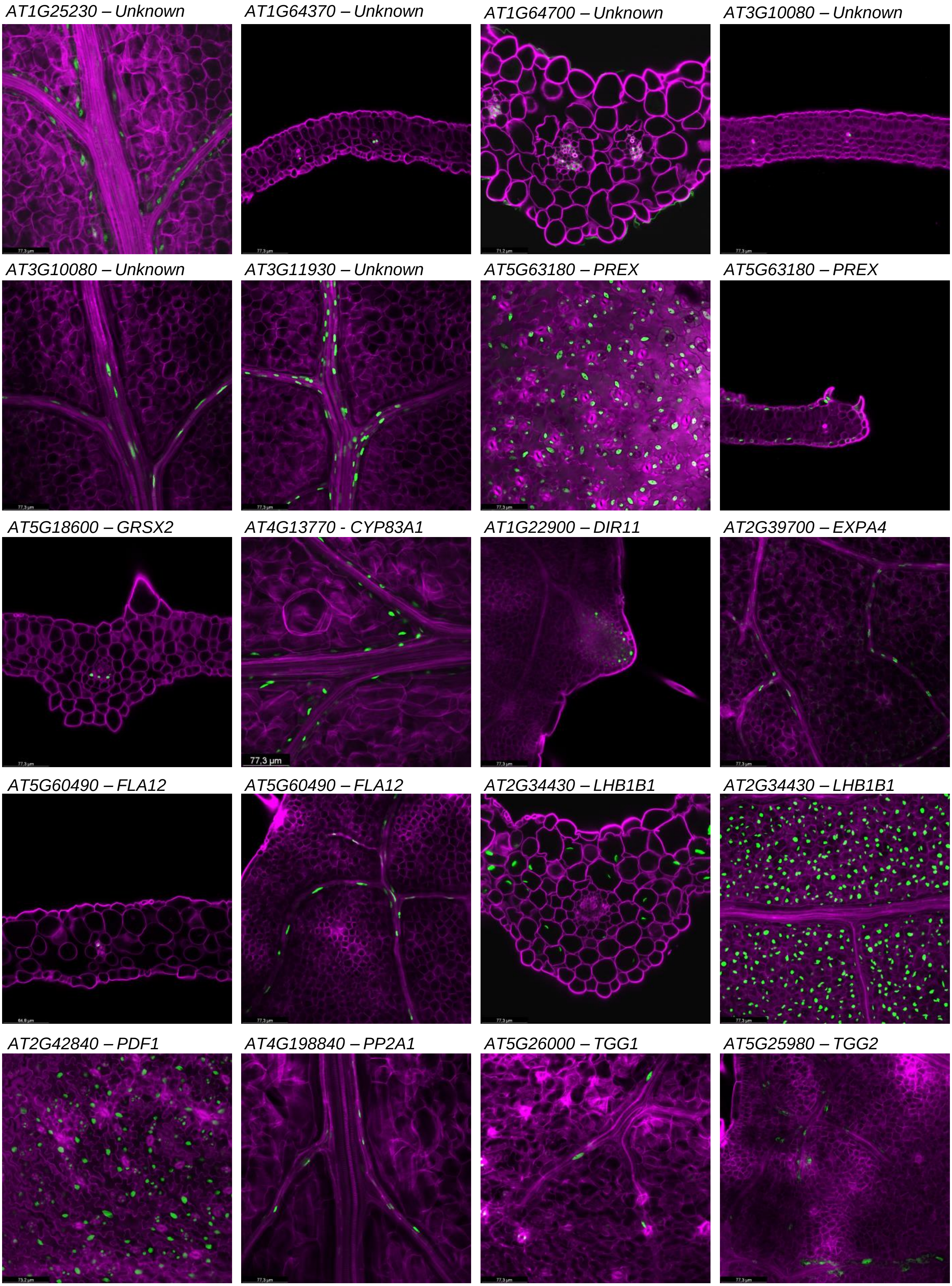


**Supp. Figure 14. Additional confocal microscopy images for the reporter lines of each tissue.** Top-view images and leaf sections are shown.

A

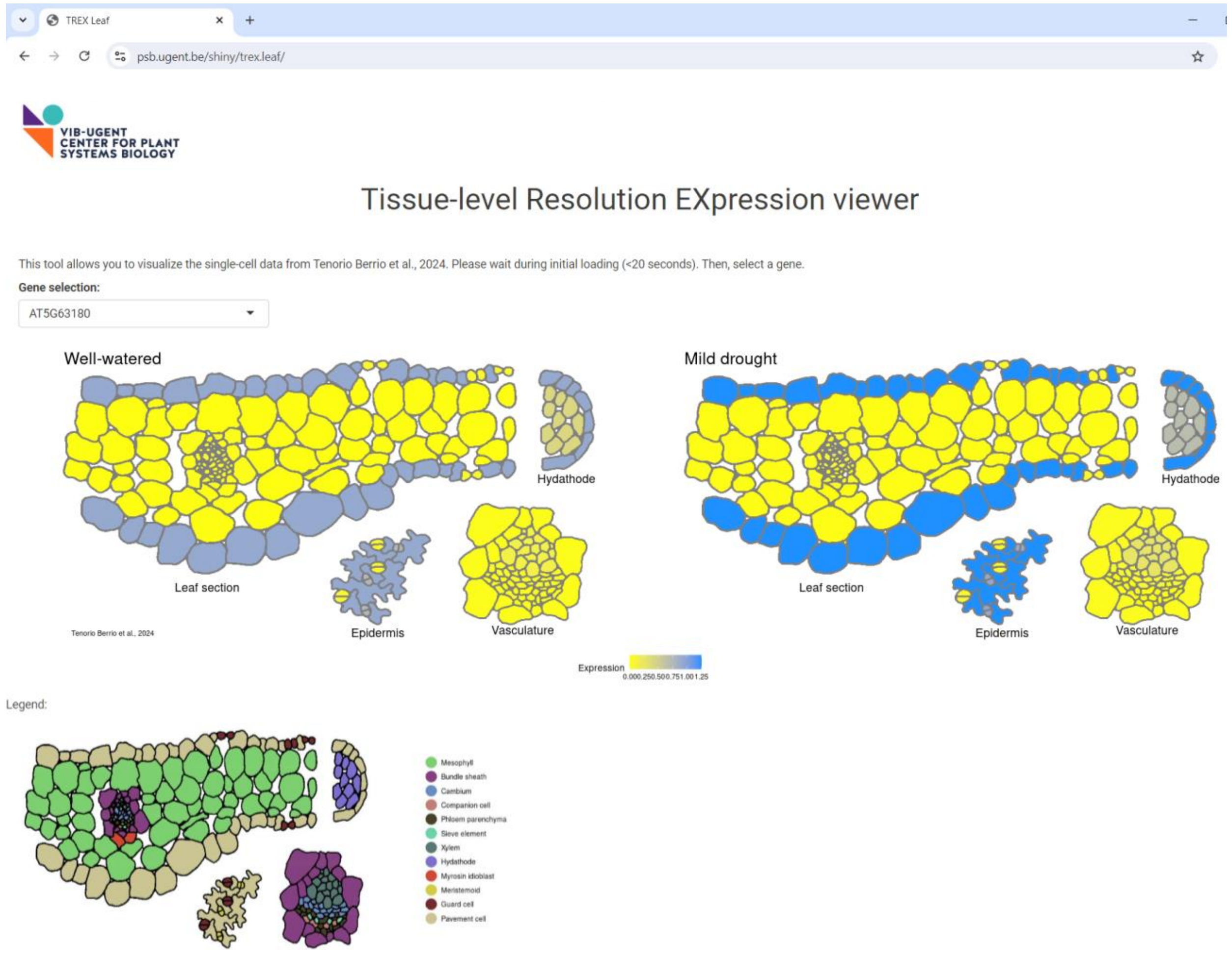


B

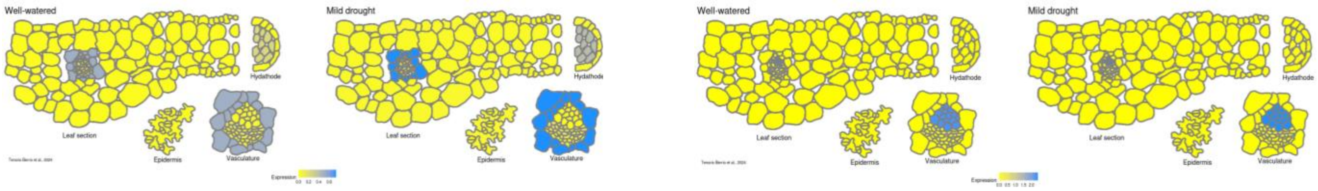


C

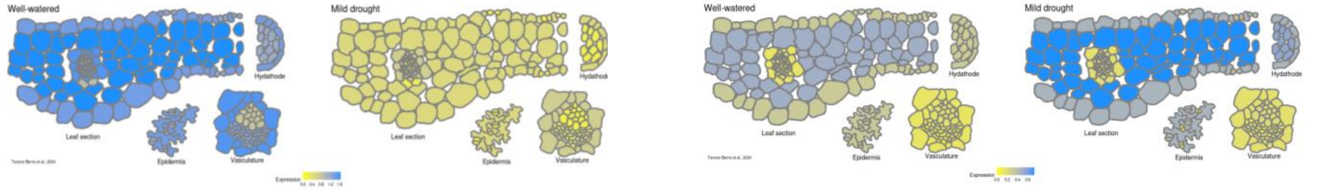


**Supp. Figure 15. Tool for single-cell data visualization in a leaf section. (A)** Screenshot of the application (https://psb.ugent.be/shiny/trex.leaf/) for visualizing the expression of any gene of interest in a schematic section of the leaf and in the epidermis. The depicted gene is the pavement cell marker *PREX* (AT5G63180). **(B)** Additional examples of tissue-specific markers *SULTR* (left) and *FLA12* (right). **(C)** Additional examples of drought-responsive genes *CSD2* (left) and *AFL1* (right).

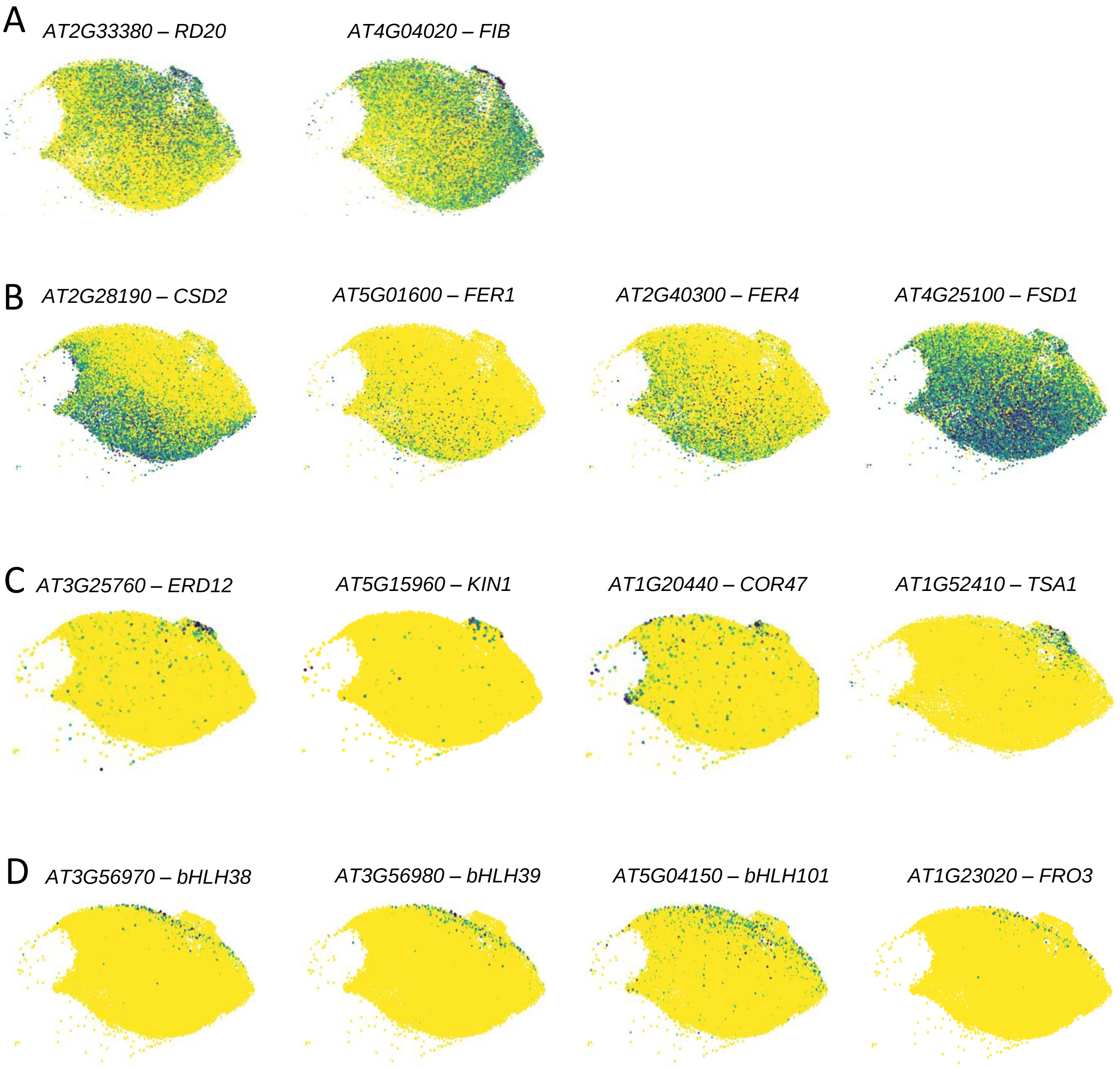


**Supp. Figure 16. Expression of drought-related genes in the mesophyll tissue. (A)** UMAP plots depicting the normalized expression of the drought induced genes *RD20* and *FIB* following the drought stress gradient. **(B)** UMAP plots depicting the normalized expression of *CSD2* and iron-related genes with decreasing expression along the drought stress gradient (described in Figure 2F). **(C)** UMAP plots depicting the normalized expression of genes expressed in the very tip of the drought gradient. **(D)** UMAP plots depicting the normalized expression of genes related to iron starvation.

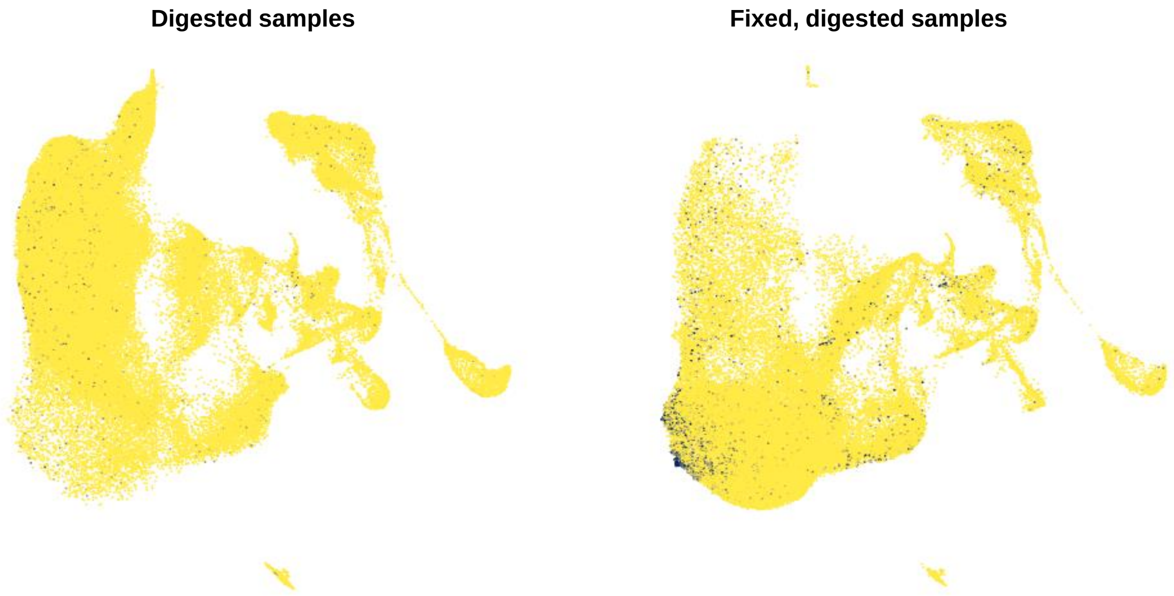


**Supp. Figure 17. Expression of *BGLU18* in the combined dataset.** UMAP plots depicting the normalized expression of *BGLU18* in the combined dataset, split by cell isolation method.

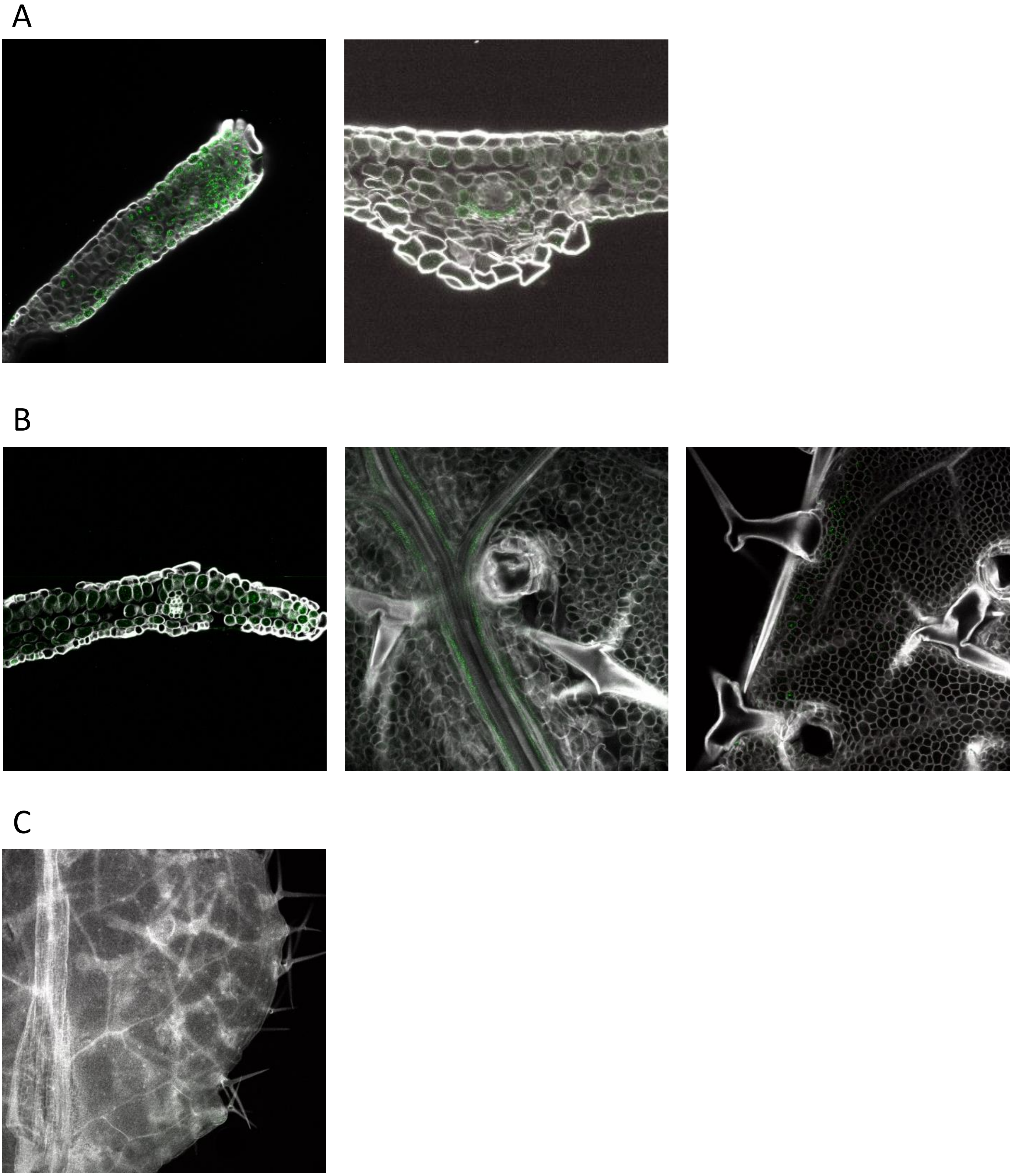


**Supp. Figure 18. Transcript visualization of canonical drought responses.** Confocal microscopy of whole-mount fluorescence *in situ* hybridization (HCR-FISH) targeting the *BGLU18* **(A)**, *TSA1* **(B)** transcripts and a negative control **(C)** upon mild drought conditions.

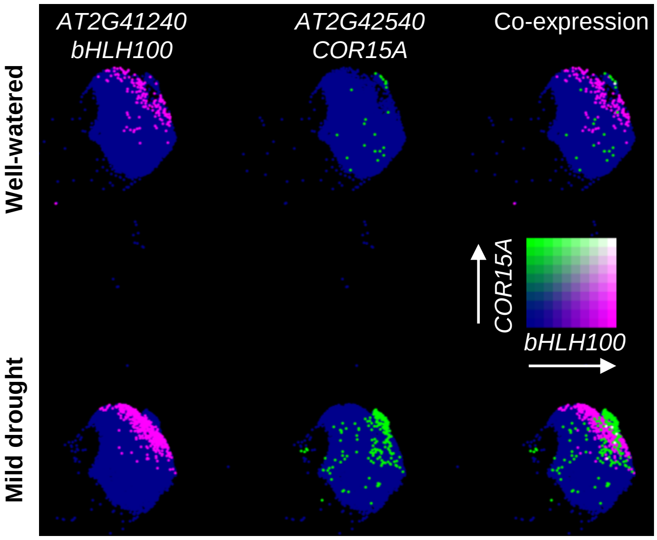


**Supp. Figure 19. Co-expression of dual drought responses in the mesophyll.** UMAP plot depicting the expression of *bHLH100* and *COR15A* in the well-watered and mild drought samples in the mesophyll.

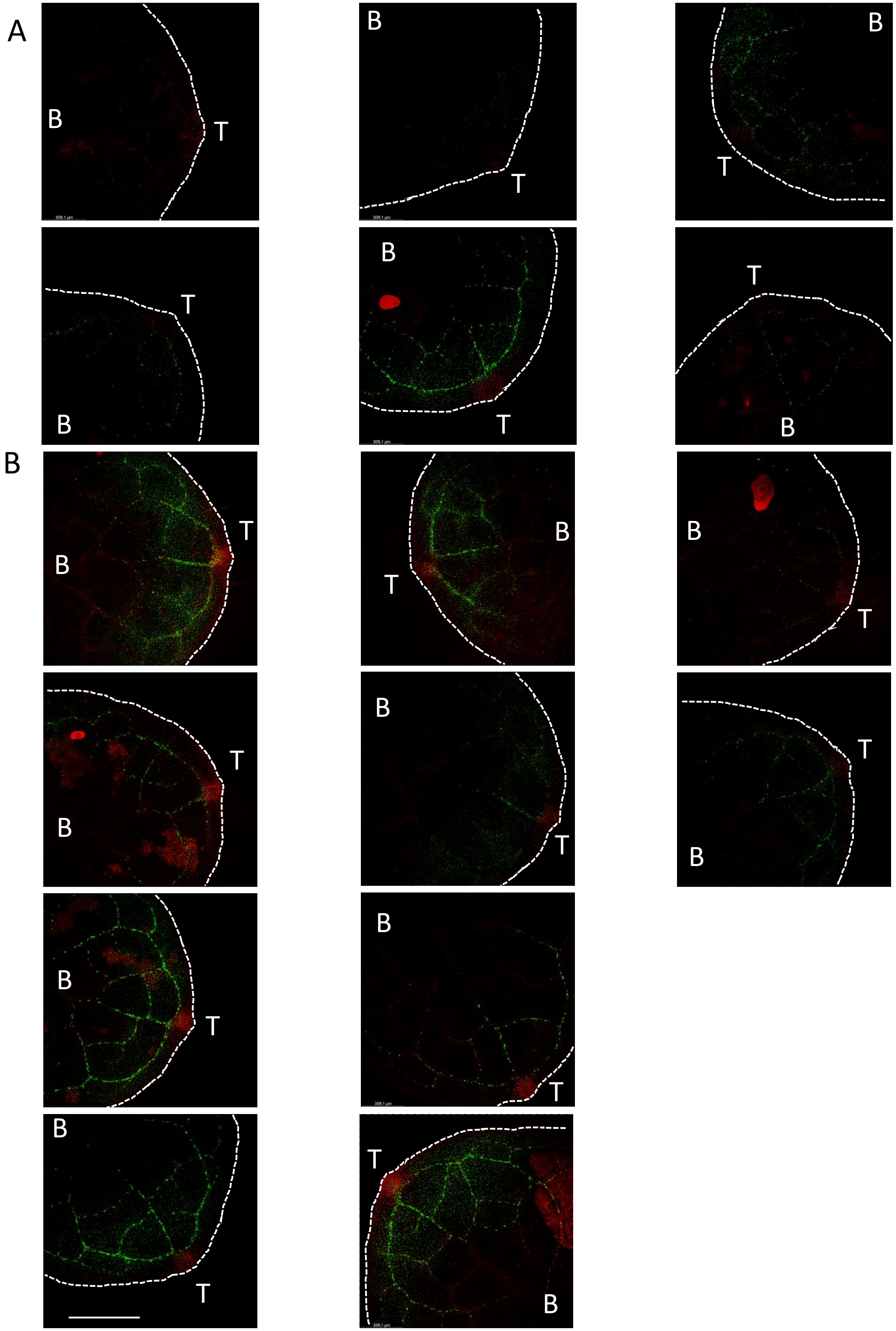


**Supp. Figure 20. Microscopy images used for image analysis of the dual drought response.** Top-view images of p*bHLH100*::nls-GFP x p*COR15A*::nls-mCherry leaves upon well-watered **(A)** or mild drought **(B)** conditions. Dashed lines delineating the margins of the leaf. T and B marking the position of the tip and the base of the leaf, respectively.
